# Prior experience establishes a proactive cortical filter that blocks redundant associations

**DOI:** 10.64898/2026.07.06.736748

**Authors:** Yoshiki Ito, Yoshihito Saito, Yusuke Atsumi, Yasuhiro Oisi, Chie Matsubara, Shigeki Kato, Kazuto Kobayashi, Kenta Kobayashi, Yumiko Yoshimura, Masanori Murayama

## Abstract

Prior experience allows selective learning by filtering out redundant cues, as exemplified by blocking, in which prior learning about one cue prevents a novel but redundant cue from being learned during subsequent compound conditioning. Top-down cortico-cortical pathways have been implicated in sensory filtering, but whether, how, and when they contribute to blocking remain unknown. Here we establish a head-fixed blocking paradigm in mice in which prior auditory cue-reward learning prevents a redundant somatosensory cue-reward association during subsequent compound conditioning. We found that a projection from anterior cingulate cortex (ACC) to primary somatosensory cortex (S1) facilitates blocking. Prior auditory learning establishes ACC-driven inhibitory control over S1 via parvalbumin neurons, before explicit somatosensory experience, thereby suppressing burst responses to the somatosensory cue, limiting network plasticity, and promoting blocking. These findings propose a proactive cortical filter that preconfigures inhibitory sensory bias to constrain future learning about redundant cues.

**Graphical abstract:** 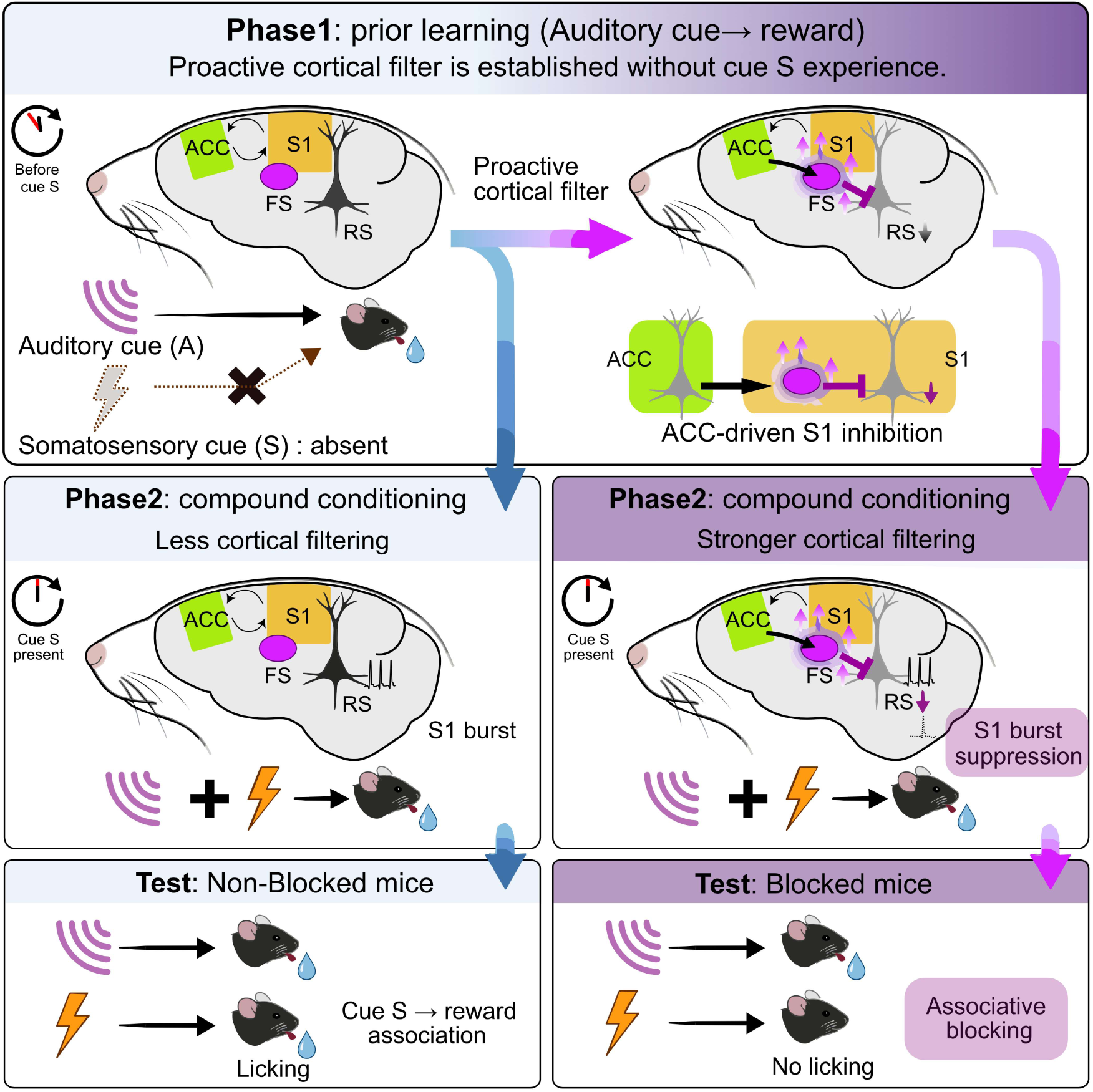

## INTRODUCTION

Animals selectively process sensory inputs by prioritizing behaviorally relevant cues and suppressing redundant ones^1–3^. Such sensory filtering is essential for adaptive perception^1,4^ and learning^5,6^, and its disruption can lead to maladaptive behaviors^7,8^, including those observed in psychiatric disorders^9,10^. In associative learning, experience-dependent filtering of redundant cue–outcome associations are classically illustrated by blocking^11^, in which prior learning about one cue (Phase1: Conditioned Stimulus 1, CS1 → Unconditioned Stimulus, US) prevents learning about a novel but redundant cue (CS2→US) during subsequent compound conditioning (Phase2: CS1+CS2→US) (Figure 1A).

**Figure 1.**
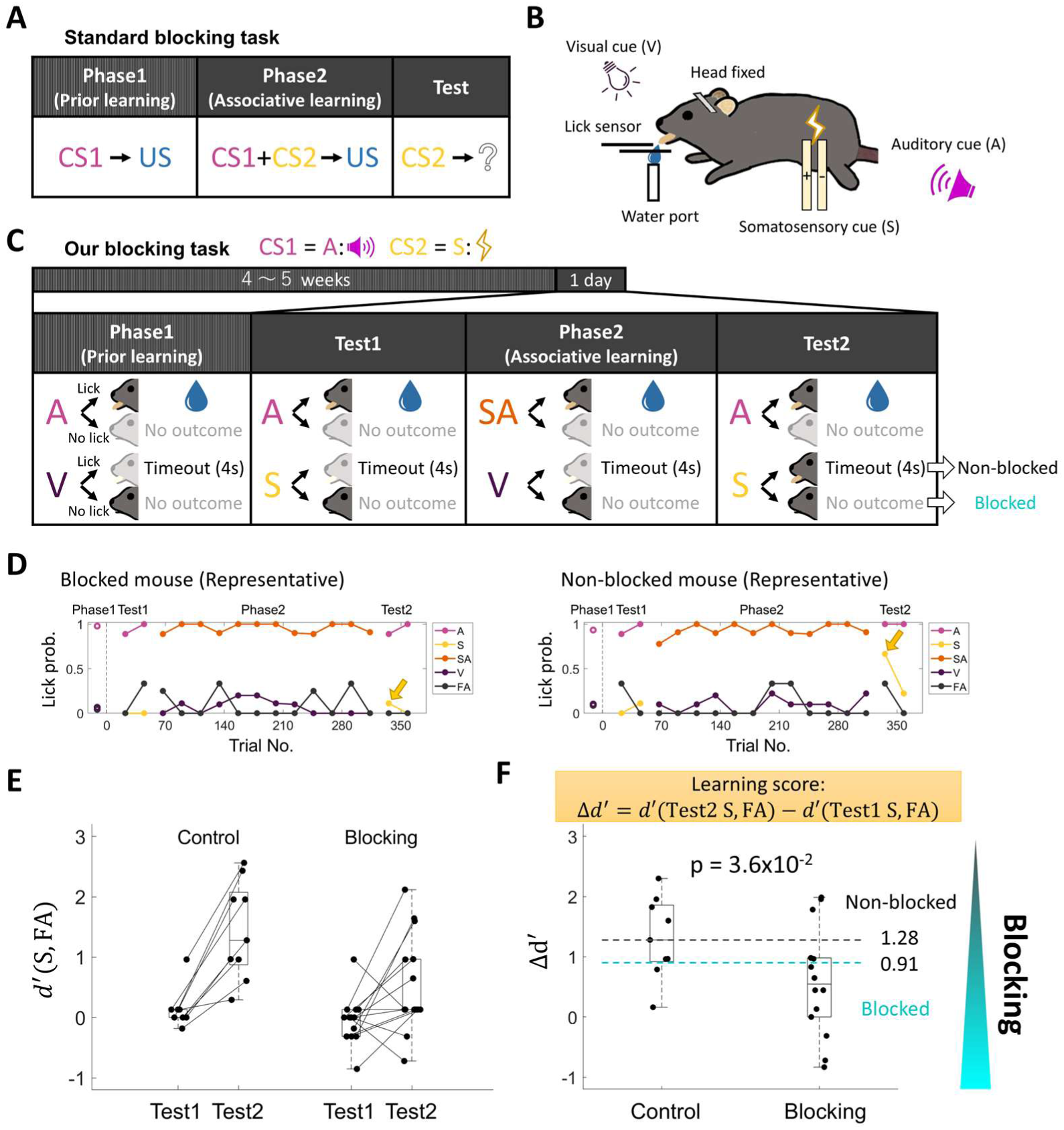
A head-fixed blocking task for quantifying inter-individual differences in cue S associability. A, A standard blocking task. In test phase, the probability of a conditioned response to CS2 is measured, which is typically low. B, Experimental setup for our blocking task with head-fixed mice. C, Our blocking task uses an auditory cue (A) for CS1, a somatosensory cue (S) for CS2, and a visual cue (V; 10 Hz light flash, 400 ms) for suppressing licking responses. Solid-colored mice indicate the major behavior, whereas faded mice indicate the minor behavior. We measure changes in licking responses to cue S in Test1 and Test2 to evaluate the blocking in each mouse. D, A representative session from a Blocked (left) and a Non-blocked (right) mouse. Both mice showed low lick probability to cue S in Test1, but only the Non-blocked mouse increased the lick probability to cue S in Test2 (yellow allows). E, The discriminability (*d*^′^) in licking responses between cue S and FA (*d*^′^(S, FA)) in Test1 and Test2 for the control task (left, n=9) and the blocking task (right, n=14). F, The difference in *d*^′^ (Δ*d*^′^) was used for the learning score. Δ*d*^′^of the blocking task was significantly lower than that of the control task (Wilcoxon one-sided rank-sum test). We used the median Δ*d*^′^ value of the control task (1.28) and the midpoint between the median Δ*d*^′^ values of the control and blocking task (0.91) to define Non-blocked (n=3/14) and Blocked mice (n=9/14), respectively.

One possible mechanism for such filtering is top-down cortical control of sensory cortex. Sensory cortical responses to cues represent not only stimulus features but also internal and contextual variables such as attention^12,13^, perception^14–17^, and behavioral relevance^18,19^, and are increasingly recognized as key determinants of cue-outcome learning^20–22^. These variables are mediated in part by top-down cortico-cortical projections, which modulate sensory representations^23–27^ and contribute to sensory-guided behaviors^28–31^. In the somatosensory system, for example, projections from dorsal frontal cortex (dFC) to the primary somatosensory cortex (S1) are required for accurate tactile perception^32^ and perceptual memory formation^33,34^.

Although subcortical circuits have been implicated in blocking^35–37^, whether top-down cortico-cortical pathways also contribute to the experience-dependent filtering of redundant associative learning remains unknown. If such pathways do contribute, they may provide a cortical mechanism that controls the specificity of cue–outcome learning by biasing credit assignment away from the redundant cue^5,38^. It also remains unclear whether this top-down mechanism is engaged reactively after redundant information is encountered or can instead proactively establish an internal bias based on prior experience before the redundant cue appears. To address these questions, we established a novel blocking paradigm with head-fixed mice in which prior learning about an auditory cue (CS1) prevents subsequent learning about a redundant somatosensory cue (CS2). Using in vivo recordings and causal manipulations, we show that prior auditory learning proactively establishes a top-down inhibitory bias in S1 before the somatosensory cue is experienced, and that this inhibitory bias promotes blocking of a redundant somatosensory association.

## RESULTS

### A head-fixed blocking task resolves individual variability in blocking

To elucidate the top-down cortical mechanisms underlying blocking, precise assessment of neural activity during behavior is required. Previous studies of blocking^11,36,39^ had two major limitations: they were performed mainly in freely moving animals and relied on group-level comparisons between CS1-conditioned and non-CS1-conditioned animals, precluding precise cue-aligned analyses and confounding neural correlates of blocking with group-dependent behavioral differences^36^. To address these issues, we established a novel blocking paradigm in head-fixed mice that allows quantitative evaluation of blocking in individual animals (Figures 1B and 1C). The task consisted of four sequential stages: Phase1 (prior learning), Test1, Phase2 (compound conditioning), and Test2. An auditory cue (A; 12,000 Hz tone, 55 dB, 150 ms) was used as CS1, and a somatosensory cue (S; 3 mA, 0.1 ms single-pulse electrical stimulation to the dorsal trunk) was used as CS2. In Phase1, mice learned the cue A-reward association. In Test1, cue S was presented alone to confirm that it did not evoke reward expectation (see STAR Methods). In Phase2, the compound cue SA was paired with reward, and in Test2, changes in licking responses to cue S were assessed. Phase1 lasted 4–5 weeks, whereas Test1, Phase2, and Test2 were performed within a single session.

In an experience-independent control task, in which a different auditory cue (A′) was used in Phase1 and Test1 (see STAR Methods), mice showed increased lick probabilities to cue S as well as to cue A in Test2 (Extended Data Figures S1A–S1D). Because cue A′ was not part of the reinforced compound cue in Phase2, this control isolated the effect of prior learning from nonspecific training effects. In contrast, in the blocking task, some mice showed low lick probabilities to cue S in Test2, indicating blocked behavior, whereas others showed high lick probabilities, indicating non-blocked behavior (Figure 1D). Note that licking responses declined in the later part of Test2 because cue S was not rewarded during the test. To quantify the cue S–reward association in Test1 and Test2, we used the *d*′ measure, defined as the difference between the lick probability to cue S and that to catch trials (i.e., false alarms; FA) (Figure 1E). The change in *d*^′^ (Δ*d*^′^ = *d*′(Test2 S, FA) − *d*′(Test1 S, FA)) was used as the learning score for somatosensory associative learning during Phase2. Mice performing the blocking task showed variable but significantly lower Δ*d*^′^ values than mice performing the control task, indicating that most animals exhibited experience-dependent suppression of novel associative learning (Figure 1F).

Rather than comparing animals performing different task conditions, we leveraged inter-individual variability within the blocking task to identify neural correlates of blocking more directly. To increase such variability, we modulated cue A intensity within the blocking task (Figure S1E). Accordingly, we defined “Non-Blocked” and “Blocked” mice using the control-task distribution of Δ*d*^′^. Mice with Δ*d*^′^ greater than the median Δ*d*^′^ of the control task (Δ*d*^′^ > 1.28) were classified as Non-Blocked, whereas those with Δ*d*^′^ lower than the midpoint between the median Δ*d*^′^ of the control and blocking tasks ( Δ*d*^′^ < 0.91) were classified as Blocked. These operational classifications were used for visualization and descriptive purposes, and are intended to reflect continuous inter-individual variability in blocking strength. This variability in Δ*d*^′^ was unlikely to reflect individual differences in reward prediction error, because Δ*d*^′^ did not significantly correlate with the number of cue S–reward pairings during Phase2, the cue A–reward or cue S–reward associations in Test1, or the learning speed of the cue A–reward association in Phase1 (Figures S1F–S1J). These results instead suggest that variability in blocking more likely reflects differences in the associability assigned to cue S.

### ACC→S1 top-down input promotes blocking

Because cue S was delivered to the dorsal trunk, we focused on the corresponding trunk region of primary somatosensory cortex (S1Tr; hereafter S1). Anatomical tracing experiments revealed reciprocal connectivity between ACC and S1: among dFC regions, the anterior cingulate cortex (ACC) directly projected to S1, with axon terminals distributed in both the upper and lower layers, consistent with a top-down cortical projection pattern^32,40^ (Figures 2A, 2B, and S2). We therefore focused on the ACC→S1 pathway in the present study.

**Figure 2.**
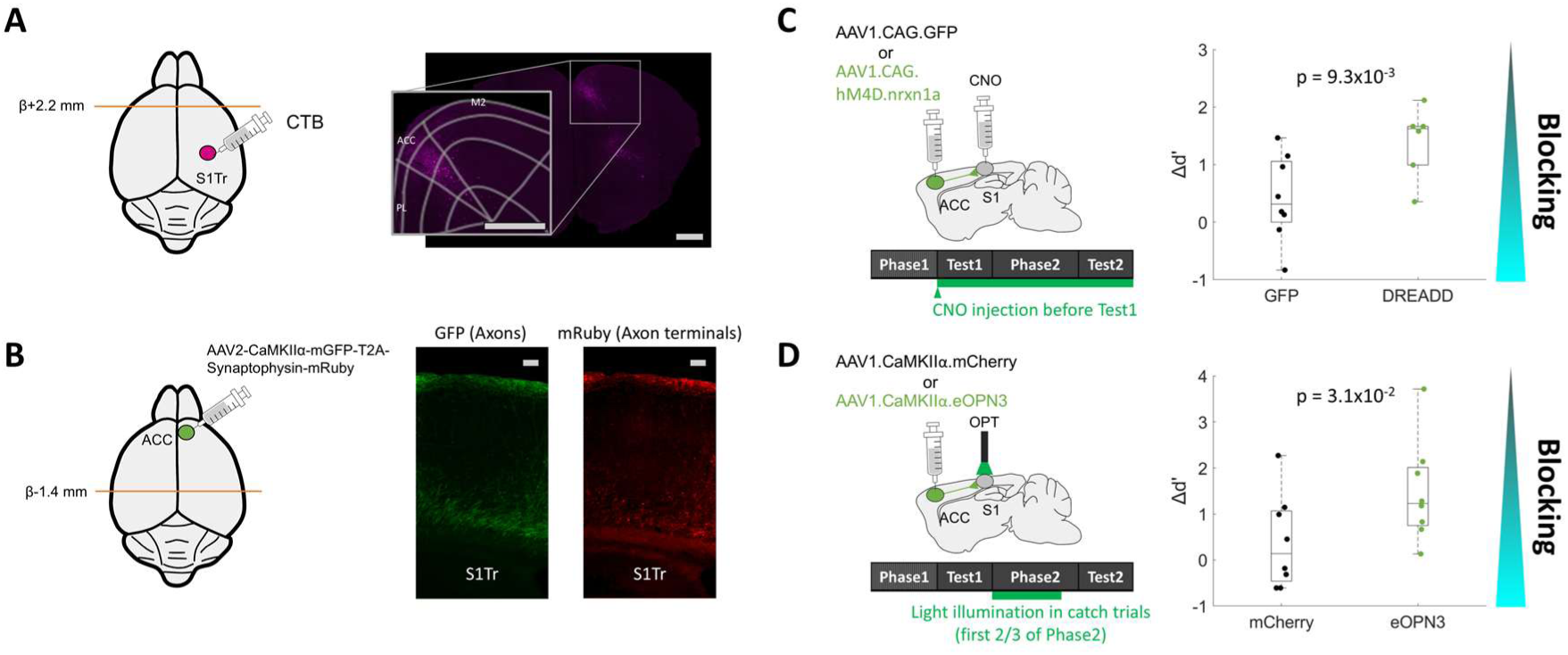
Top-down projection from ACC to S1Tr facilitates blocking. A, A retrograde tracer (CTB) was injected into S1Tr. Labeled neurons were clustered at approximately 2.2 mm anterior to bregma. Based on anatomical criteria, these populations were identified as the anterior cingulate cortex (ACC). Scale bar, 500 μ m. B, To visualize axons and presynaptic terminals from ACC to S1Tr with different fluorescent markers (GFP and mRuby), AAV2-CaMKII α-mGFP-T2A-Synaptophysin-mRuby was injected into ACC. Axons and axon terminals from ACC were observed in both the superficial and deep layers of S1Tr. Scale bar, 100 μ m. C, Silencing of ACC terminals at S1 with inhibitory DREADD during Test1, Phase2 and Test2 significantly increased Δ*d*^′^compared with the GFP control group (Wilcoxon one-sided rank-sum test, n=8 for control and n=6 for DREADD). D, Optogenetic silencing of ACC terminals at S1 with eOPN3 during Phase2 significantly increased Δ*d*^′^ compared with the mCherry control group (Wilcoxon one-sided rank-sum test, n=8 each).

Because dFC→S1 input has been reported to be essential for accurate somatosensory perception^32^ and its perceptual memory^33,34^, we initially expected that suppressing this top-down input during somatosensory associative learning would attenuate somatosensory perception and thereby facilitate blocking. To test this, we first expressed inhibitory designer receptors exclusively activated by designer drugs (DREADDs) in ACC and suppressed ACC axon terminals from Test1 to Test2 by locally injecting clozapine-N-oxide (CNO) into S1. Contrary to our expectation, suppression of ACC terminals significantly increased the learning score (Δ*d*^′^) compared with the GFP control (Figure 2C). To further support this result, we next used optogenetics to selectively suppress ACC axonal activity in S1 during Phase2, delivering photostimulation during catch trials in the first two-thirds of Phase2. This manipulation also significantly increased the learning score (Δ*d*^′^) relative to control mice (Figure 2D). Thus, contrary to our initial expectation, we found that ACC→S1 top-down input facilitates blocking.

### ACC top-down input drives S1 FS units early in compound conditioning

To investigate the circuit mechanism, we next simultaneously recorded neural activity in ACC and S1 across Test1, Phase1, and Test2, classifying units as regular-spiking (RS, putative excitatory neurons) or fast-spiking (FS, putative parvalbumin, PV, neurons (Figures 3A and B). Cue S trials in Test1 were termed Test1 S, and cue SA trials in Phase2 were divided into six periods (SA-1 to SA-6), each consisting of approximately 20 trials.

**Figure 3.**
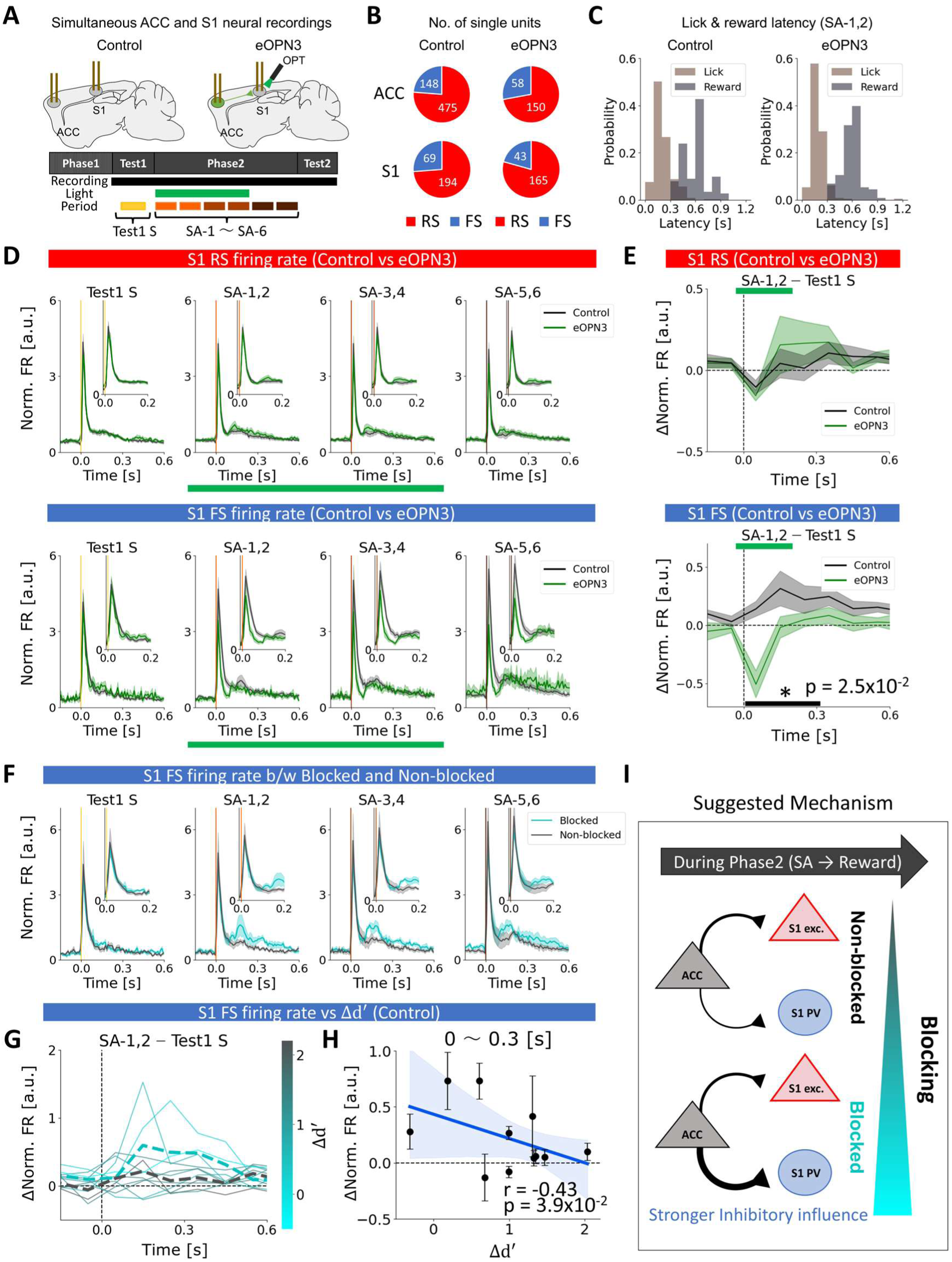
Top-down input increases S1 FS firing rate. A, Unit activity in ACC and S1 was simultaneously recorded during Test1, Phase2 and Test2 from untreated control mice and eOPN3 mice using silicon probes. Light illumination of eOPN3 mice was delivered to eOPN3 mice during the first two-thirds of catch trials in Phase2. B, Total numbers of RS and FS units from control mice (n=11) and eOPN3 mice (n=7). C, Histogram of lick and reward latencies of all control mice and eOPN3 mice in SA-1,2. D, Mean normalized firing rates (FR) of all S1 RS (Top) and S1 FS (Bottom) units from control (gray) and eOPN3 (green) mice in each period. Firing rates were normalized by the mean firing rate within 600 ms after cue onset in Test1 S. Mean ± s.e.m. Green bars represent light illumination periods in eOPN3 mice. E, Differences in normalized firing rates between SA-1,2 and Test1 S for S1 RS (Top) and S1 FS (Bottom). eOPN3 mice (green) showed a significantly lower S1 FS firing rate for 300 ms after cue onset compared with control mice (gray) (One-sided permutation test with max-statistic correction across time). F, Mean normalized firing rates of S1 FS units of Blocked (cyan) and Non-blocked (black) control mice in each period. Mean ± s.e.m. G, Differences in normalized S1 FS firing rates between SA-1,2 and Test1 S. Solid lines represent individual animals colored by Δ*d*^′^, and dashed lines indicate the group mean of Blocked (cyan) and Non-blocked (black) control mice. H, Changes in normalized S1 FS firing rates within 300 ms after cue between SA-1,2 and Test1 S were plotted against Δ*d*^′^. A significant negative correlation was observed (one-sided bootstrap test). I, Suggested mechanism in which Blocked mice exhibit stronger top-down input influence on S1 PV neurons than Non-blocked mice.

For each epoch, we calculated the firing rates of S1 RS and S1 FS units and normalized them to the mean firing rate during Test1 S trials to align inter-individual variability in unit recordings. After presentation of the compound cue SA during Phase2, licking behavior typically began within 300 ms, and reward was delivered 300 ms after cue onset in both control and eOPN3 mice (Figure 3C). Whereas the cue-evoked firing rates of S1 RS units did not differ markedly between control and eOPN3 mice, those of S1 FS units were reduced in eOPN3 mice after top-down input suppression (Figure 3D). Importantly, the decrease in S1 FS activity emerged from the early Phase2 trials (SA-1,2), in which cue SA elicited significantly lower S1 FS firing rate during the first 300 ms in eOPN3 mice than in control mice, whereas no such difference was observed in S1 RS units (Figure 3E). This reduction in S1 FS firing in eOPN3 mice was unlikely to be fully explained by differences in lick probability (Figures S3A), lick latency (Figures S3B), or ACC RS firing rates (Figures S4A and S4B). Together, these results suggest that top-down input from ACC selectively drives S1 FS units from the earliest phase of compound conditioning.

We next asked whether this early increase in S1 FS firing was related to the strength of blocking. Blocked mice tended to show higher cue-evoked S1 FS firing rates after cue SA than Non-Blocked mice (Figure 3F). Notably, the increase in cue-evoked firing rate within the first 300 ms after cue SA in SA-1,2, relative to cue S in Test1 S, was greater in Blocked mice and showed significant negative correlations with the learning score (Δ*d*^′^) (Figures 3G and 3H). This blocking-related increase in S1 FS firing was also unlikely to be fully explained by differences in lick probability (Figure S3C), lick latency (Figure S3D), or ACC RS firing rates (Figures S4C and S4D). Together, these results suggest that stronger ACC-driven recruitment of S1 PV neurons during the earliest phase of compound conditioning contributes to blocking (Figure 3I).

### ACC-driven PV recruitment suppresses S1 burst responses linked to blocking

Because overall firing rates of S1 RS units changed little despite pronounced recruitment of S1 FS units, we next asked whether top-down inhibition alters the temporal structure rather than the mean firing rate of RS spiking. We therefore examined burst firing in S1 RS units, because burst activity has been implicated in perception^32^ and perceptual learning^33^. Comparing Test1 S and SA-1 revealed that majority of S1 RS units in Blocked mice showed a reduction in proportion of burst firing in SA-1 relative to Test1 S (Figure 4A). To quantify this effect, we calculated the proportion of burst events relative to all firing events in each neuron and defined the burst index^41^ (BI, see STAR Methods) (Figures 4B and 4C).

**Figure 4.**
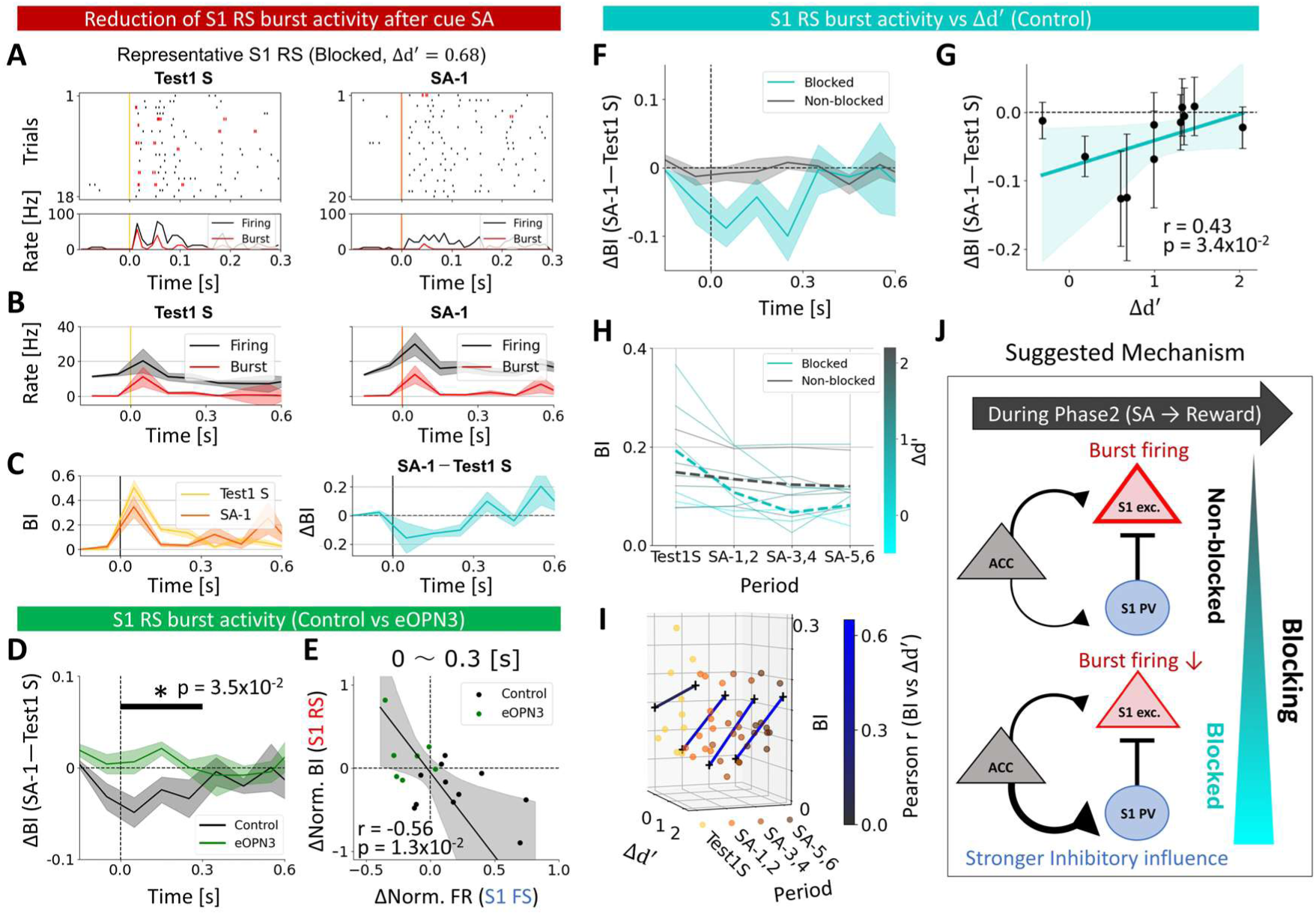
Top-down input suppresses S1 RS burst activity in Blocked mice. A, Raster (Top) and mean event rates (Bottom) of a representative S1 RS unit from a Blocked mouse (black, firing event; red, burst event) in Test1 S (Left) and SA-1 (Right). B, Mean firing and burst event (100 ms bins) of all S1 RS units in a Blocked mouse in Test1 S (Left) and SA-1 (Right). Mean ± s.e.m. C, (Left) Time course of burst index (BI; 100 ms bins) in Test1 S (yellow) and SA-1 (orange) for all S1 RS units in a Blocked mice. (Right) The difference in BI between SA-1 and Test1 S. Mean ± s.e.m. D, Time course of the difference in BI between SA-1 and Test1 S in control (gray) and eOPN3 mice (green). They significantly differed for 300 ms after cue onset. E, The differences in S1 FS normalized firing rates between SA-1 and Test1 S within 300 ms after cue were plotted against the differences in BI between SA-1 and Test1 S within 300 ms after cue. A significant negative correlation was observed (one-sided bootstrap test). Black dots represent control mice, and green dots represent OPN3 mice. F, The differences in BI between SA-1 and Test1 S of Blocked (cyan) and Non-blocked (gray) control mice. Mean ± s.e.m. G, The differences in BI between SA-1 and Test1 S of control mice for 300 ms after cue were plotted against Δ*d*^′^. They showed a significantly positive correlation (one-sided bootstrap test). Error bars, Mean ± s.e.m. Mean regression (cyan) ± 95% CI (shaded). H, BI within 300 ms after cue onset at each period. Solid lines, individual animals (colored by Δ*d*^′^); dashed lines, group mean. I, Linear regression analysis between BI and Δ*d*^′^across each period, colored by correlation coefficients. J, Suggested mechanism in which strong inhibitory influence on S1, particularly in Blocked mice, suppresses burst responses of S1 excitatory neurons to somatosensory input and facilitates blocking.

In control mice, BI in SA-1 decreased following cue onset compared with that in Test1 S. In contrast, this decrease was not observed in eOPN3 mice, and a significant difference was found between control and eOPN3 mice within 300 ms after cue onset (Figure 4D). These results suggest that suppression of burst firing in RS units during the early stages of Phase2 depends on ACC→S1 top-down input. Anatomical analyses^42^ showed that PV neurons are the major inhibitory neuronal population receiving ACC projections (Figure S5), suggesting that this burst suppression is mediated by S1 PV neurons. We therefore tested whether the change in BI from Test1 S to SA-1 was related to the change in cue-evoked firing rates of S1 FS units and found a significant negative correlation (Figure 4E). In contrast, the differences in cue-evoked firing rates of S1 RS units did not show a significant correlation with those of S1 FS units (Figure S6A). Together, these results suggest that top-down input from ACC, from the very earliest phase of Phase2, drives S1 PV neurons and thereby selectively suppresses burst firing in excitatory neurons without substantially altering their overall firing rates.

Given that ACC top-down input suppressed S1 RS burst firing from the earliest phase of Phase2, we next asked whether this early reduction in S1 burst responses was related to the strength of blocking. Blocked mice tended to show a greater BI reduction in SA-1 relative to Test1 S than Non-Blocked mice (Figure 4F), and this reduction was significantly correlated with the learning score (Δ*d*^′^) (Figure 4G). Moreover, in Blocked mice, BI in response to cue SA progressively decreased over the course of Phase2 (Figure 4H), and BI showed a significant positive correlation with Δ*d*^′^ during late Phase2 (Figures 4I and S6B). Together, these results suggest that early suppression of S1 burst responses by ACC top-down input contributes to blocking, most likely through recruitment of S1 PV neurons.

### Pre-existing ACC–S1 inhibitory bias predicts Blocked versus Non-blocked behaviors

Given the ACC-mediated increase in S1 PV activity and decrease in S1 RS bursting, we next looked at interactions between these regions across learning, by calculating functional connectivity (Figure 5A and STAR Methods). Functional connectivity between ACC and S1 RS showed only a weak negative correlation with the learning score (Δ*d*^′^) in Test1 S, but this negative correlation gradually strengthened over the course of Phase2 (from SA-1,2 to SA-5,6) (Figure 5B). In contrast, functional connectivity between ACC and S1 FS already showed a strong negative correlation with Δ*d*^′^in Test1 S, and this correlation structure was maintained throughout Phase2 (Figures 5C–5D). These findings suggest that inter-individual differences in the strength of top-down inhibitory influence on S1 were already present before Phase2 and might subsequently shape both ACC–S1 RS network plasticity and learning outcome. To test this idea, we examined the relationship between functional connectivity bias in Test1 S, defined as the difference between ACC–S1 RS and ACC–S1 FS functional connectivity, and network plasticity, defined as the absolute change in ACC–S1 RS functional connectivity between SA-1,2 and SA-5,6. These measures showed a significant positive correlation (Figure 5E). Furthermore, functional connectivity bias in Test1 S was also positively correlated with Δ*d*^′^, and this relationship was altered by top-down inhibition during Phase2 (Figures 5F and S7).

**Figure 5.**
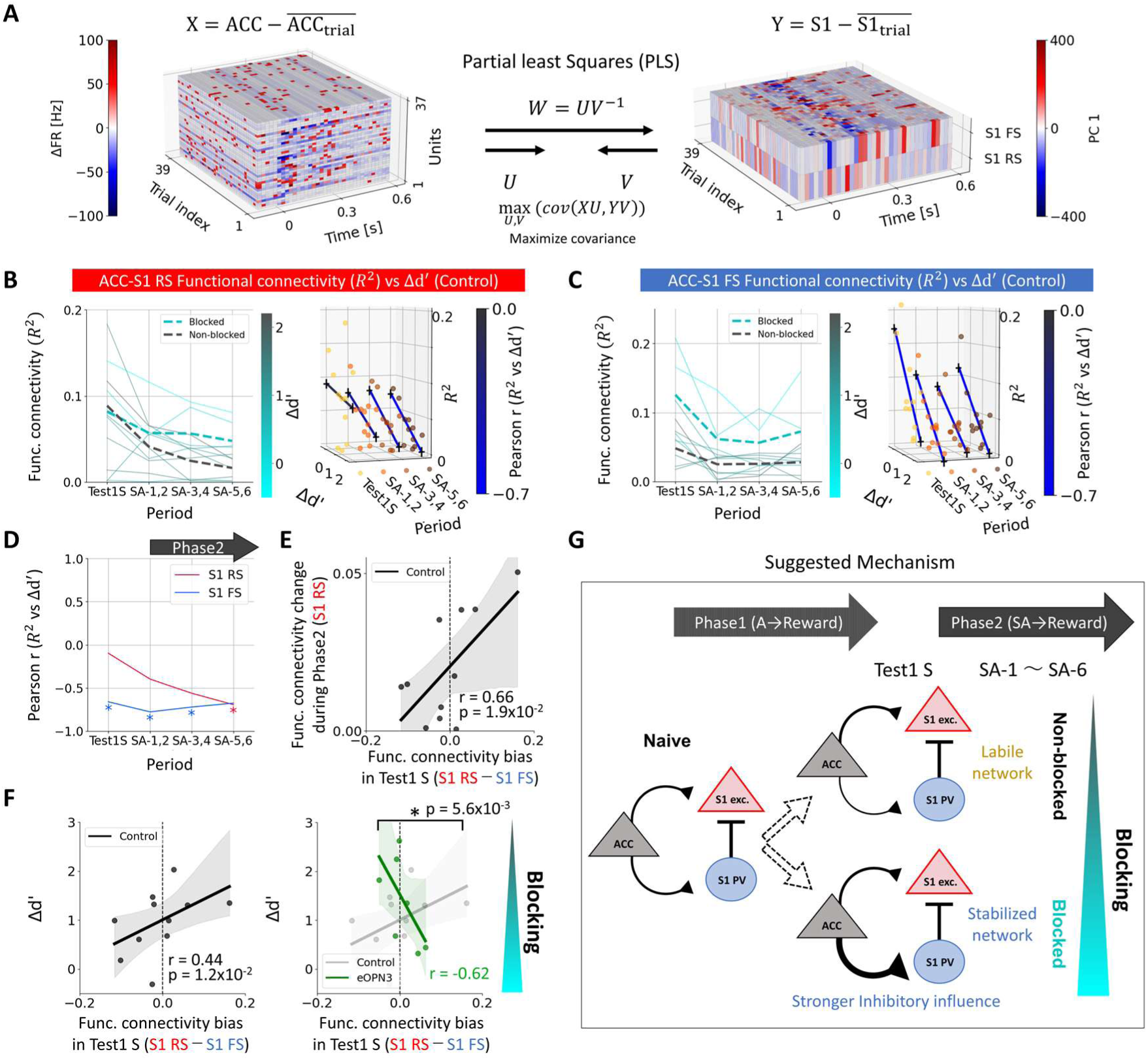
ACC-S1 functional connectivity bias before learning explains subsequent learning performance. A, Schematic of estimating ACC–S1 functional connectivity using partial least squares (PLS). B, (Left) Functional connectivity (R²) between ACC and S1 RS across each period. Solid lines indicate individual animals colored by Δ*d*^′^, and dashed lines indicate the group mean. (Right) Linear regression analysis between ACC-S1 RS functional connectivity and Δ*d*^′^ across each period, colored by correlation coefficients. C, Same analysis as B for ACC-S1 FS functional connectivity. D, Correlation coefficients between ACC-S1 RS or ACC-S1 FS functional connectivity and Δ*d*^′^ across each period. Significant negative correlations were observed for ACC–S1 FS functional connectivity during Test1 S (*P* = 4.4*x*10^-2^), SA-1,2 (*P* = 2.7*x*10^-2^), and SA-3,4 (*P* = 2.2*x*10^-2^), and for ACC–S1 RS functional connectivity during SA-5,6 (*P* = 4.1*x*10^-2^) (*P < 0.05; one-sided bootstrap test). E, Functional connectivity bias (*R^2^_S1 RS_* − *R^2^_S1 FS_*) in Test1 S was plotted against absolute ACC–S1 RS functional connectivity changes during Phase2 (*R^2^_S1 RS_(SA-5,6)* − *R^2^_S1 FS_(SA-1,2)|*) in control mice. They showed a significant positive correlation (one-sided bootstrap test). Mean regression ± 95% CI (shaded). F, (Left) Functional connectivity bias (*R^2^_S1 RS_* − *R^2^_S1 FS_*) in Test1 S was plotted against Δ*d*^′^ in control mice. They showed a significant positive correlation (one-sided bootstrap test). (Right) The same analysis in eOPN3 mice. The correlation coefficient was significantly different from control mice (two-sided label permutation test). Mean regression ± 95% CI (shaded). G, Suggested mechanism in which individual differences in top-down inhibitory influence on S1 before Phase2 predicts the result of somatosensory associative learning in Phase2.

Together, these results suggest that Blocked mice start Phase2 with a stabilized ACC–S1 network state, shaped by a stronger inhibitory influence from ACC to S1, which suppresses subsequent novel associative learning of somatosensory stimuli. In contrast, Non-Blocked mice start Phase2 with a more labile network state, in which weaker inhibitory influence permits greater ACC–S1 RS plasticity and facilitates learning (Figure 5G). These findings raised the possibility that proactive cortical filtering is already established before compound conditioning begins, despite the absence of an explicit somatosensory cue during Phase1.

### Non-somatosensory prior learning strengthens ACC-driven S1 inhibition before explicit somatosensory experience

Because the preceding analyses suggested that inhibitory bias differs across animals before Phase2 begins, we next tested whether prior non-somatosensory learning during Phase1 establishes this difference in top-down influence on S1. To do so, we used fiber photometry to monitor S1 excitatory or PV neuronal responses to optogenetic activation of ACC in head-fixed mice at rest before and after Phase1^31^ (Figure 6A and 6B). ChRmine^43^ was expressed in ACC, and GCaMP8m was expressed either in putative excitatory neurons using the CaMKIIα promoter or in PV neurons in S1. Red light pulses were delivered to ACC, and ACC-evoked Ca²⁺ responses were recorded from the S1 cortical surface (Figures 6C, S8A–S8C).

**Figure 6.**
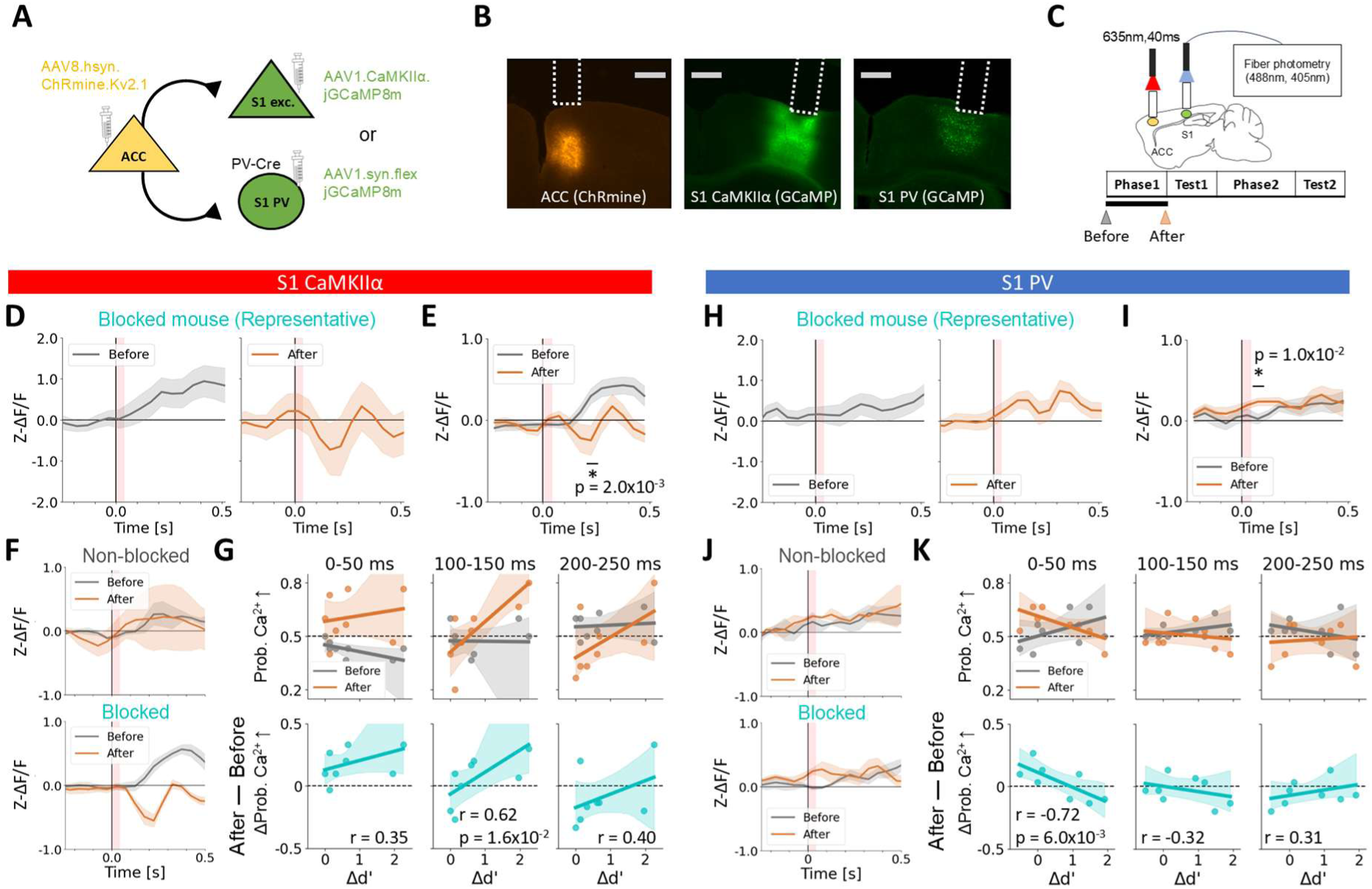
Blocked mice establish ACC-driven S1 inhibition during non-somatosensory learning. A, ChRmine was expressed in ACC, and GCaMP was expressed in S1 excitatory (exc.) neurons in WT mice (CaMKII α) or S1 PV neurons in PV-Cre mice (syn-flex). B, Representative sections showing expression of ChRmine in ACC and GCaMP in S1. Scale bar, 500 μ m. Dotted lines indicate estimated fiber positions. C, ACC-activated S1 Ca^2+^ responses were recorded using multi-channel fiber photometry (488 nm for GCaMP; 405 nm for artifact correction) before (gray) and after (orange) Phase1. D, Mean S1 exc. Ca^2+^ responses of a representative Blocked mice before (Left) and after (Right) Phase1. Mean ± s.e.m. Red shaded areas indicate the ACC activation window (40 ms). E, Mean S1 exc. Ca^2+^ responses of all recorded mice (n=8) before and after Phase1. Mean ± s.e.m. Red shaded areas indicate the ACC activation window (40 ms). S1 exc. Ca^2+^ responses was significantly reduced after Phase1 (*P < 0.05; one-sided paired bootstrap test with max-T correction). F, Mean S1 exc. Ca^2+^ responses of Blocked (top, n=6/8) and Non-blocked (bottom, n=2/8) mice. Mean ± s.e.m. The negative S1 exc. Ca^2+^ responses were prominent in Non-blocked mice. G, (Top) The probability of trials showing an increase in S1 exc. Ca^2+^ responses relative to 50 ms preceding ACC activation onset was quantified at 0-50 (Left), 100-150 (Middle), and 200-250 (Right) ms after the onset before (gray) and after (orange) Phase1. (Bottom) Changes in S1 exc. Ca^2+^ response probability from before to after Phase1 were plotted against Δ*d*^′^. The significant positive correlation was observed at 100-150 ms (one-sided bootstrap test). Mean regression (solid line) ± 95% bootstrap confidence interval (CI, shaded). H-K Same analysis as D-G for S1 PV Ca^2+^ responses (n=8; Blocked mice, n=4/8; Non-blocked mice, n=2/8). The significant negative correlation was observed at 0-50 ms (one-sided bootstrap test).

In home-caged control mice, ACC-evoked Ca²⁺ responses did not differ over time in either S1 excitatory or S1 PV neurons (Figure S8D). By contrast, in mice performing the blocking task, ACC-evoked responses in S1 excitatory neurons were significantly reduced after Phase1 (Figures 6D and 6E), whereas responses in S1 PV neurons were significantly increased (Figures 6H and 6I). Both changes were more pronounced in Blocked than in Non-Blocked mice (Figures 6F and 6J), and the corresponding changes in response probability were significantly correlated with the learning score (Δ*d*^′^) (Figures 6G and 6K). Notably, these effects developed progressively as task difficulty increased during Phase1 by introducing a visual No-Go cue and reducing the auditory cue intensity to enable one-session learning (Figures S9A–S9E). Together, these results indicate that non-somatosensory associative learning progressively strengthens ACC-driven inhibition onto S1 via local PV neurons, particularly in animals that later exhibit blocking. Because no explicit somatosensory cue was presented during Phase1, this learning-dependent shift is more consistent with contextual experience than with direct sensory experience.

## DISCUSSION

Overall, we identify a previously unrecognized mechanism that we term a proactive cortical filter, in which experience establishes top-down inhibitory control over a task-irrelevant sensory cortex before explicit presentation of redundant sensory information (Figure 7). This filter emerged as a preconfigured ACC–S1 network state characterized by enhanced recruitment of S1 PV neurons (Figures 3E-3H), suppression of burst responses to sensory cues (Figures 4D-4I), and inhibitory functional connectivity bias (Figures 5E and 5F). Animals with a well-established proactive cortical filter during Phase1 showed redundant associative blocking (Figures 6G, and 6K), and suppression of ACC→S1 top-down input abolished this effect and allowed the associative learning (Figures 2C and 2D). Together, these findings highlight the experience-dependent preconfiguration of cortical top-down filter that gates future associative learning.

**Figure 7.**
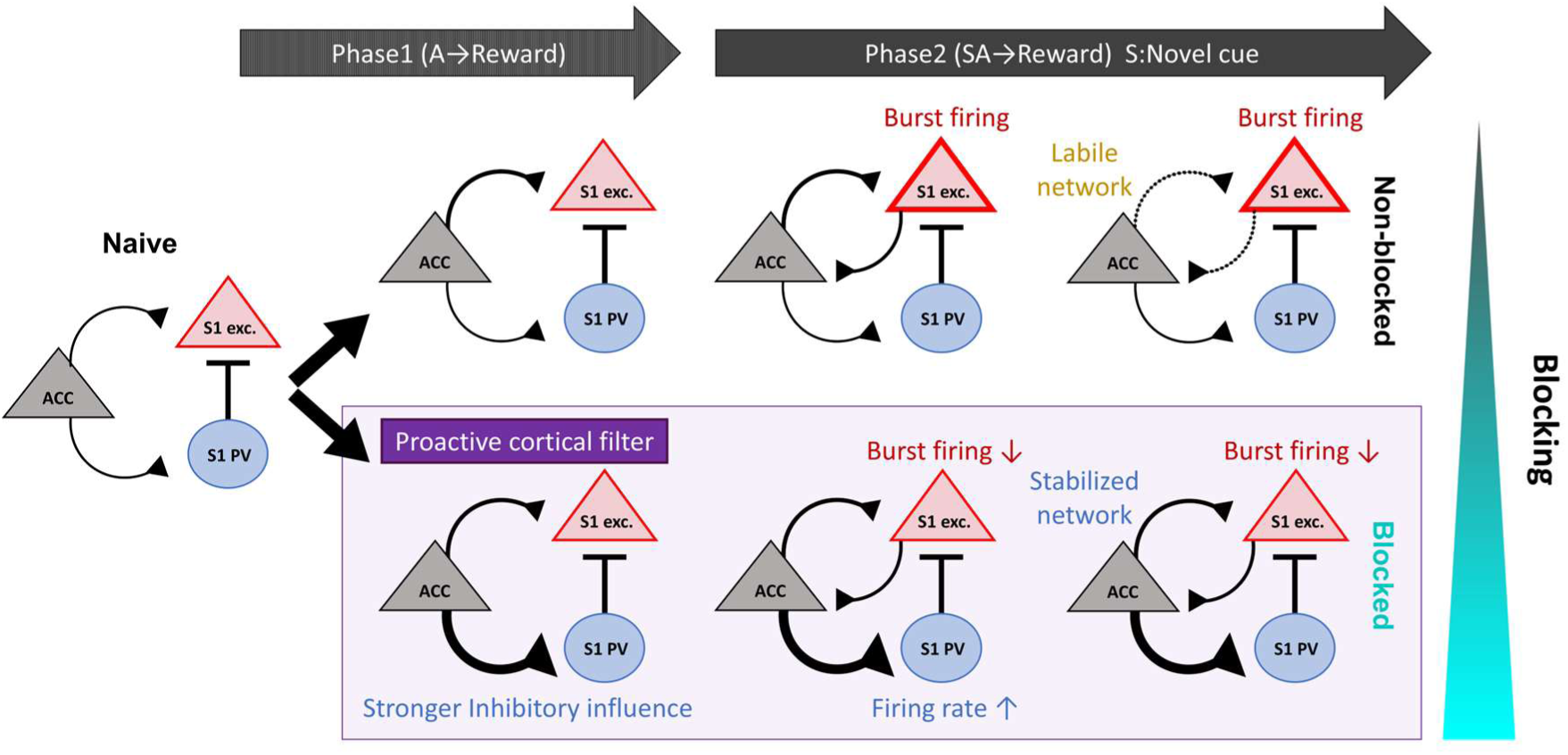
Schematic summary of the proactive cortical filter facilitating blocking. Line thickness indicates estimated strength of synaptic connections. In S1 excitatory (exc.) neurons, box outline thickness represents burst responses to cue S. The proactive cortical filter refers to a mechanism whereby contextual experience alone strengthens top-down inhibitory influence on S1 via S1 PV neurons. This proactive cortical filter suppresses burst responses of S1 excitatory neurons to novel somatosensory inputs during learning and prevents subsequent associative learning.

Previous studies of cortical filtering have emphasized the reactive mechanisms of suppressing ongoing sensory inputs, such as self-generated or predictable signals, by top-down inhibitory control of sensory cortex via local PV neurons^23,44,45^. The key distinction from these previous studies is that cortical filtering in our study was established proactively, before the redundant sensory cue was ever presented, thereby exerting its function in future associative blocking. Notably, ACC-driven inhibition of S1 increased during Phase1 and became more pronounced as task difficulty increased (Figure S9), raising the possibility that the attentional demand on processing task-relevant cues contributed to the establishment of this filter. Such a proactive filter may confer at least four physiological advantages. It may minimize distraction by novel cues, preserve established knowledge, prevent plasticity from being wasted on redundant cues, and reduce the metabolic cost imposed by these processes^46^.

The proactive filter identified here may represent one source of inter-individual variation in learning ability, depending on whether it is formed and how strongly it is expressed. Previous works have emphasized inter-individual variability in associative learning, focusing on attentional processes^47,48^, cue selectivity^49^, and related neural correlates^50,51^. Our findings extend these theoretical and correlative accounts by identifying that proactive cortico-cortical mechanisms causally contribute to the variability in learning. Given the prominent role of sensory filtering deficits including associative blocking in some neuropsychiatric disorders^9,10^, together with the broader relevance of sensory processing abnormalities^52,53^, understanding individual differences in proactive cortical filtering may also provide insight into maladaptive learning in these disorders.

Our findings also extend classical views of blocking. Blocking has long been interpreted in relation to prediction error signals in subcortical reinforcement systems^35,54^. Our results suggest that blocking can be regulated not only by global reinforcement signals such as dopamine^36,39^, but also by cue-specific top-down cortical mechanisms. Notably, ACC input suppressed S1 activity before reward delivery (Figures 3C-3E), making it unlikely that this effect simply reflects reward prediction error itself. Rather, the ACC→S1 pathway may provide a circuit mechanism through which prior learning selectively reduces the associability of a redundant sensory cue before that cue acquires substantial associative strength. This interpretation is more closely aligned with attentional theories of associative learning^5,6,55^, which propose that associability is regulated in a cue-specific manner, increasing for informative cues and decreasing for redundant sensory cues.

One possible mechanism underlying this reduced associability is proactive filtering-mediated suppression of burst activity in sensory cortex. Burst firing has been implicated in perception^16,56^, learning^29,57^, and effective cortical output^58,59^, and our findings suggest that reduced burst firing contributes to blocking. Importantly, burst responses to cue S were selectively suppressed when the same somatosensory stimulus was presented as part of cue SA, despite little change in mean firing rate (Figures 4E and S5A). This observation raises the possibility that top-down control of burst firing in sensory cortex contributes to cue-specific credit assignment^38,60,61^. Prior learning may establish an experience-dependent inhibitory bias in the ACC→S1 pathway, which subsequently suppresses sensory cortical burst responses to the redundant sensory cue before reward delivery, thereby biasing credit assignment away from that cue. At least two circuit-level implementations could, in principle, give rise to such burst suppression: excitatory top-down influence onto S1, which is known to support accurate tactile perception^32^ and perceptual memory^33^, could be reduced, or excitatory top-down input could drive local inhibitory circuits^23,44,45^. Our results are more consistent with the latter implementation in the context of blocking.

Future work should examine the synaptic mechanisms of proactive cortical filtering. One possible mechanism is anti-Hebbian plasticity at ACC synapses onto S1 PV neurons^62,63^. Because S1 is largely inactive while ACC neurons are repeatedly engaged by the auditory cue-reward associative learning, this mismatch in pre- and postsynaptic activity may progressively strengthen ACC recruitment of PV neurons, thereby establishing the proactive cortical filter. If anti-Hebbian plasticity indeed underlies the proactive aspect of sensory filtering, variability in innate projection patterns, synaptic plasticity, and receptor expression within this cortico-cortical circuit may account for the inter-individual differences in, and dysfunction of, selective learning.

### Limitations of the study

First, in the electrophysiological experiments, untreated mice were used as controls for eOPN3-expressing mice. Second, although our loss-of-function experiments established the necessity of ACC→S1 top-down inputs for proactive cortical filtering, the mixed excitatory and inhibitory influences of ACC projections on S1 precluded a complementary gain-of-function test for blocking enhancement.

## RESOURCE AVAILABILITY

The data and code that support the findings of this study are available from the corresponding author upon reasonable request.

## Lead contact

Further information and requests for materials and resources should be directed to and will be fulfilled by the lead contact, Masanori Murayama.

## Materials availability

This study did not generate new, unique reagents.

## Data and code availability

- All data reported in this paper will be shared by the lead contact upon request.
- All custom code has been deposited at https://github.com/toppo365/proactive_filter.git.
- Any additional information required to reanalyze the data reported in this paper is available from the lead contact upon request.

## ACKNOWLEDGMENTS

We thank T. Yoneda for valuable comments and discussions. This research was supported by Japan Society for the Promotion of Science KAKENHI grants (JP24KJ0566 to Y.I.; JP20H05775 and JP24H02313 to M.M.; 24K02140, Grant-in-Aid for Scientific Research (B) to Y.Y.), by AMED-Brain/Minds 1.0 project (JP15dm0207001 to M.M.) and AMED-Brain/Minds 2.0 project (JP23wm0625001 to M.M. and K.K (Kenta)), by the Cooperative Study Program of NIPS (25NIPS159 to Y.Y), by KAO Corp. (to M.M.), and by the Toray Science Foundation (to M.M.).

## AUTHOR CONTRIBUTIONS

Conceptualization, Y.I., Y.S., Y.A., and M.M..; methodology, Y.I., Y.S., Y.A., and M.M.; Investigation, Y.I..; writing – original draft, Y.I.; writing – review & editing, M.M.; funding acquisition, Y.I., K.K (Kenta), Y.Y., and M.M.; resources, Y.O., C.M. S.K., K.K. (Kazuo), and K.K. (Kenta); supervision, Y.Y., and M.M.

## Supplementary figures

**Figure S1.**
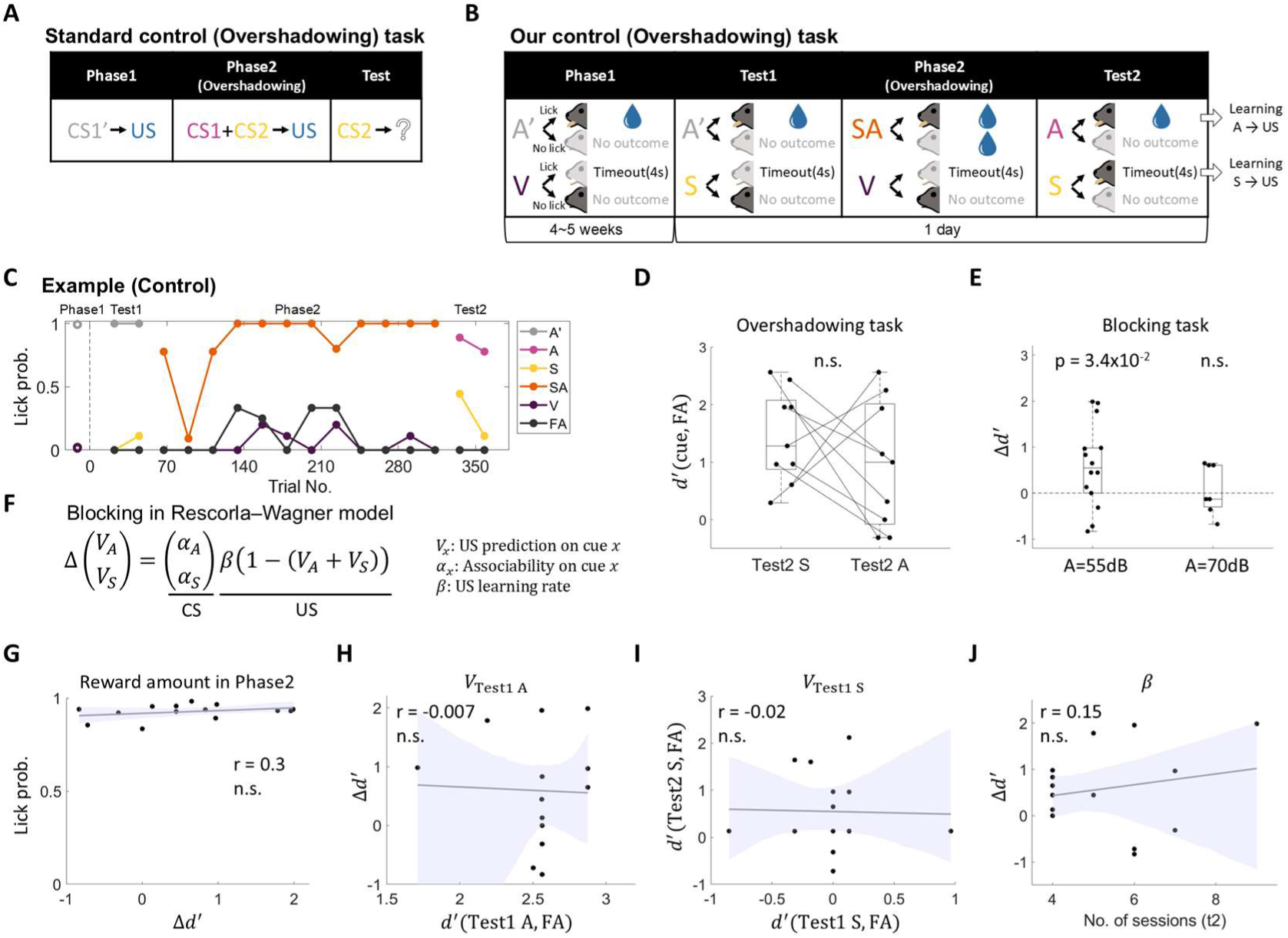
The learning score (**Δ*d***^′^) likely reflects associability assigned to cue S, related to Figure 1. A, A standard control task. In Phase1, a cue (CS1’) distinct from CS1 and CS2 is associated with US. In Phase2, novel cues (CS1 and CS2) are associated with US, allowing assessment of learning driven by cue competition between CS1 and CS2 (i.e., overshadowing). B, Our control task. A distinct auditory cue (A’) was used in Phase1 and Test1, and mice experienced paired presentations of novel cues (S and A) and reward in Phase2. Reward was provided regardless of licking responses to cue SA during Phase2 to prevent task disengagement. Solid-colored mice indicate the major behavior, whereas faded mice indicate the minor behavior. C, A representative session of the control task. This mouse showed licking responses to both Test2 A and Test2 S, indicating that cue A–reward and cue S-reward association were both learned. D, The discriminability between Test2 S and FA rate (*d*^′^(Test2 S, FA)) and between Test2 A and FA rate (*d*^′^(Test2 A, FA)) for the control task. No significant difference was observed (Wilcoxon one-sided signed-rank test, testing the hypothesis *d*^′^(Test2 S, FA) > *d*^′^(Test2 A, FA), n=9). E, The learning scores (Δ*d*^′^) when cue A intensity was changed from 55 dB (n=14) to 70 dB (n=6) in the blocking task. Whereas the median of Δ*d*^′^ when A=55dB was significantly different from zero, that when A=70 dB was not (Wilcoxon two-sided signed-rank test). F, Formal expression of blocking in a standard associative learning model (Rescorla-Wagner model^54^). Mackintosh (1975)^5^ explained blocking by reduction in cue S associability (*α*_S_) during Phase1 due to the selective attention to cue A. G, The relationship between lick probability in Phase2 and the learning score (Δ*d*^′^). H, Relationship between *d*^′^(Test1 A, FA) and the learning score (Δ*d*^′^). I, Relationship between *d*^′^(Test1 S, FA) and *d*^′^(Test2 S, FA). J, Relationship between the number of sessions required for associative learning on the 80 dB auditory cue (A) and reward in Phase1 (t2, see STAR Methods) and the learning score (Δ*d*^′^). For panels G-J, mean regression lines (solid) with 95% CI (shaded) are shown. No significant correlations were observed (bootstrap test, n.s., n=14). These results indicate that the number of S–reward pairings in Phase2, reward prediction for cue A or S after Phase1 (*V*_Test1 A_ or *V*_Test1 S_, ′espectively), or associative learning rate (*β*) did not explain Δ*d*^′^, suggesting that inter-individual variability in learning score (Δ*d*^′^) is largely attributable to variability in cue S associability (*α*_S_) in Rescorla-Wagner model^54^.

**Figure S2.**
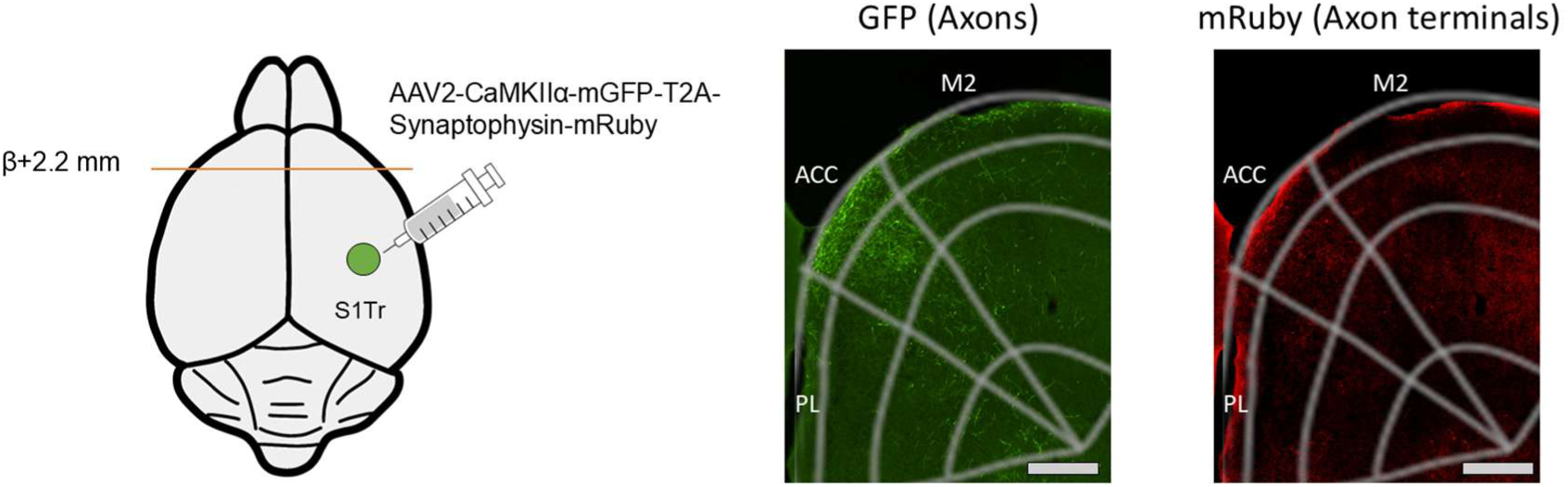
Anatomy of the S1Tr to ACC bottom-up projection, related to Figure 2. To confirm the reciprocal projection between ACC and S1Tr, axons and presynaptic terminals from S1Tr projecting to ACC were labeled with different fluorescent markers (GFP and mRuby) by injecting AAV2-CaMKII α-mGFP-T2A-Synaptophysin-mRuby into S1Tr. Axons and axon terminals from S1 were observed in ACC. Scale bar, 200 μ m.

**Figure S3.**
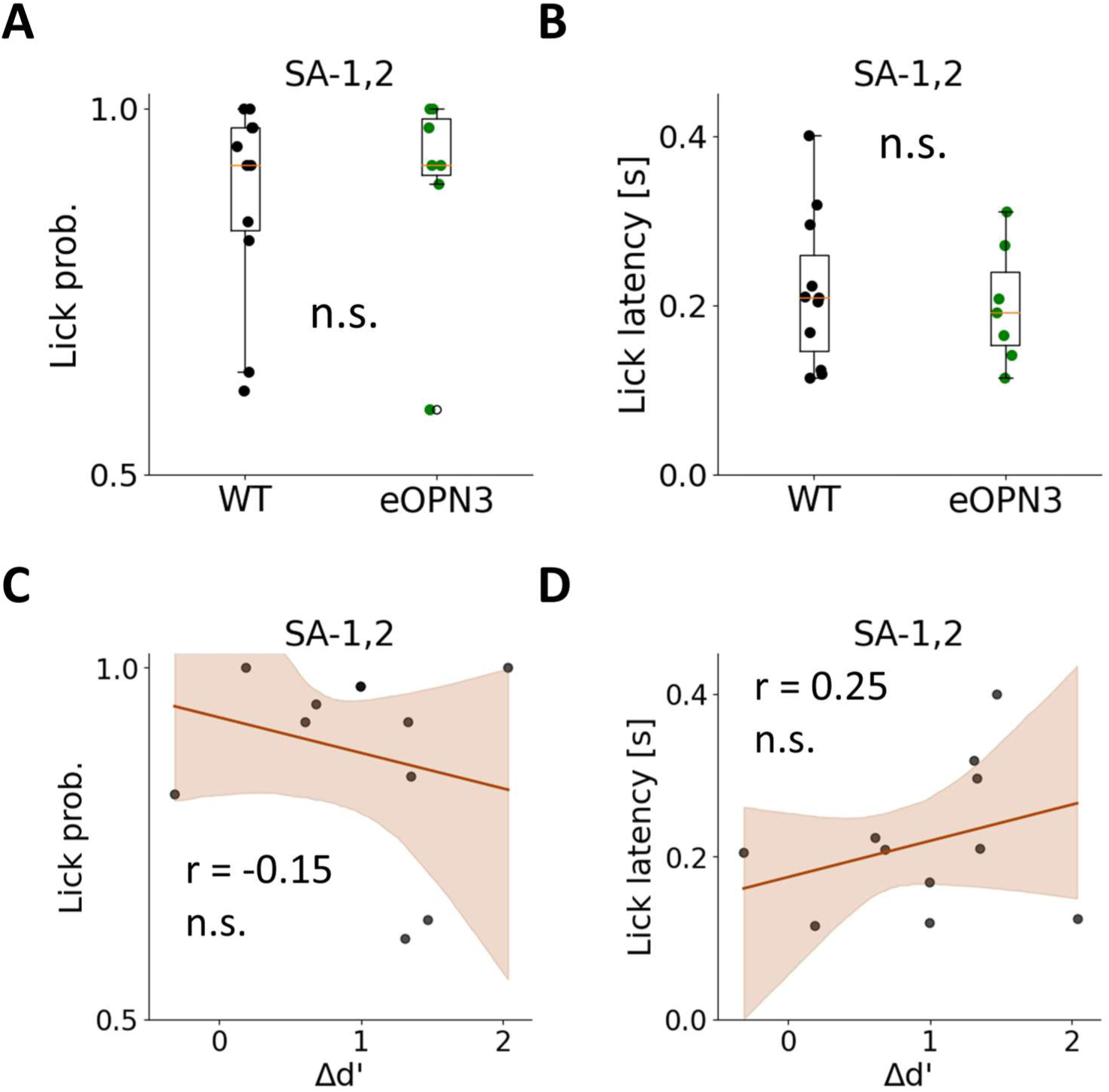
Lick probability and lick latency in SA-1,2, related to Figure 2. A, Lick probability in SA-1,2 of control (gray, n=11) and eOPN3 (green, n=7) mice. No significant difference was observed (Wilcoxon rank-sum test). B, Lick latency in SA-1,2 of control (gray, n=11) and eOPN3 (green, n=7) mice. No significant difference was observed (Wilcoxon rank-sum test). C, Lick probability in SA-1,2 of control mice was plotted against the learning score (Δ*d*^′^). Mean regression (solid line) ± 95% CI (shaded). No significant correlation was observed (bootstrap test, n.s., n=11). D, Lick latency in SA-1,2 of control mice was plotted against the learning score (Δ*d*^′^). Mean regression (solid line) ± 95% CI (shaded). No significant correlation was observed (bootstrap test, n.s., n=11).

**Figure S4.**
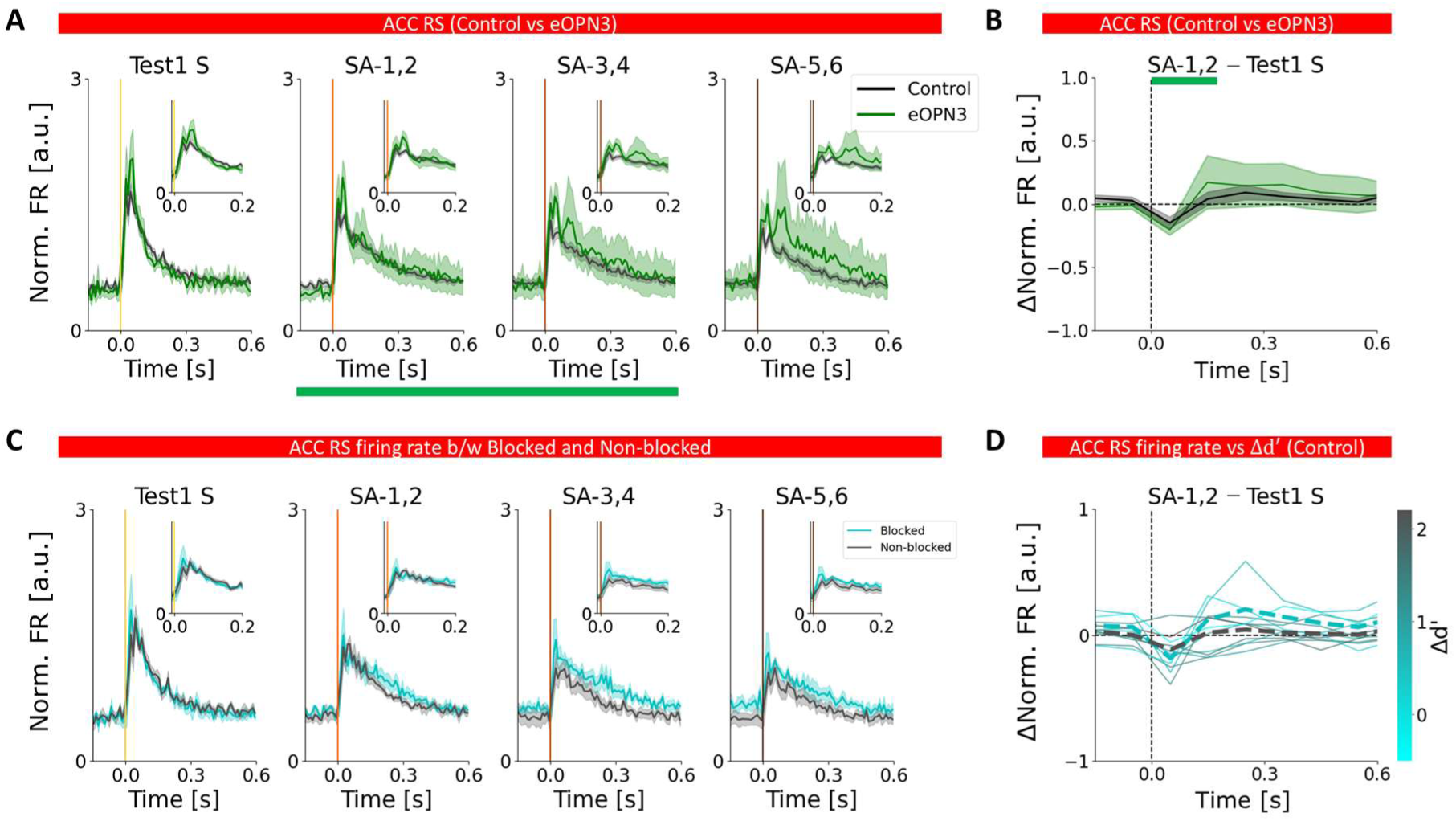
Changes in ACC RS firing rates did not majorly contribute to changes in S1 FS firing rate, related to Figure 3. A, Mean normalized firing rates of ACC RS units of control (gray, n=11) and eOPN3 (green, n=7) mice in each period. Mean± s.e.m. Insets show an expanded view of the 0–0.2 s time window. B, Differences in normalized firing rate of ACC RS between SA-1,2 and Test1 S. No prominent difference was observed. (One-sided permutation test with max-statistic correction across time) C, Mean normalized firing rates of ACC RS units of Blocked (cyan) and Non-blocked (black) control mice in each period. Mean ± s.e.m. D, Differences in normalized S1 FS firing rates between SA-1,2 and Test1 S. Solid lines represent individual animals colored by Δ*d*^′^, and dashed lines indicate the group mean of Blocked (cyan) and Non-blocked (black) control mice. Blocked mice exhibited higher ACC RS firing rates than Non-blocked mice in SA-1,2, consistent with the increased firing rates observed in S1 FS neurons. However, the increase in ACC RS firing rates was relatively modest, suggesting that the differences in ACC RS to S1 FS synaptic connectivity between Blocked and Non-blocked mice, rather than ACC RS activity itself, may play a more important role in determining the increase in S1 FS firing rate in SA-1,2.

**Figure S5.**
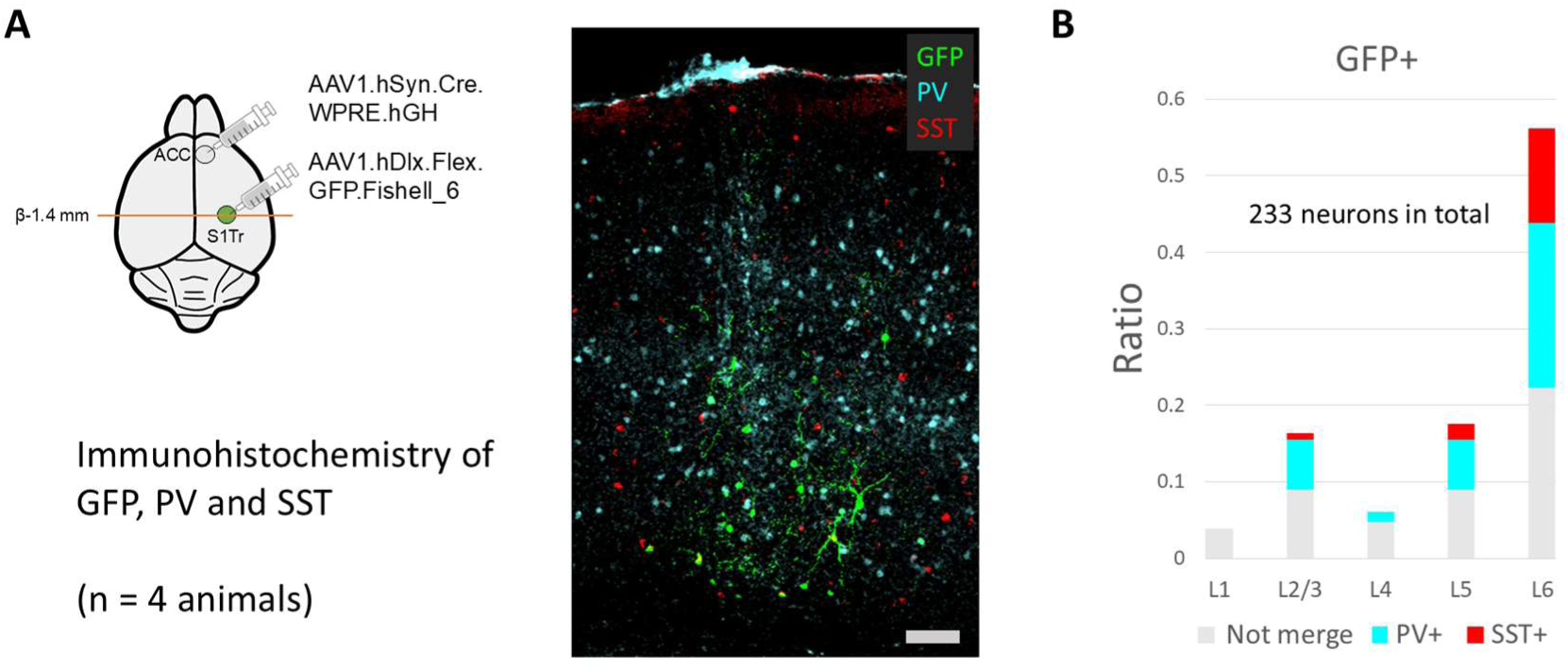
Labeling of S1Tr inhibitory neurons receiving projections from ACC, related to Figure 4. A, (Left) Labeling of S1 inhibitory neurons receiving projections from ACC using DISECT^42^. Taking advantage of the anterograde transsynaptic property of AAV1-syn-Cre virus vector, S1Tr inhibitory neurons receiving projections from ACC were labeled with GFP by injecting AAV1-hSyn-Cre in ACC and AAV1-hDlx-flex-GFP in S1Tr. (Right) Immunohistochemistry was performed to label PV-positive (cyan) and SST-positive (red) neurons, and their co-expression with GFP was quantified. Scale bar, 100 μ m. B, Proportions of PV- and SST-positive neurons among GFP-positive neurons. GFP-positive neurons were predominantly located in the deep layers of S1Tr, and among these, PV-positive neurons showed the highest percentage of co-expression.

**Figure S6.**
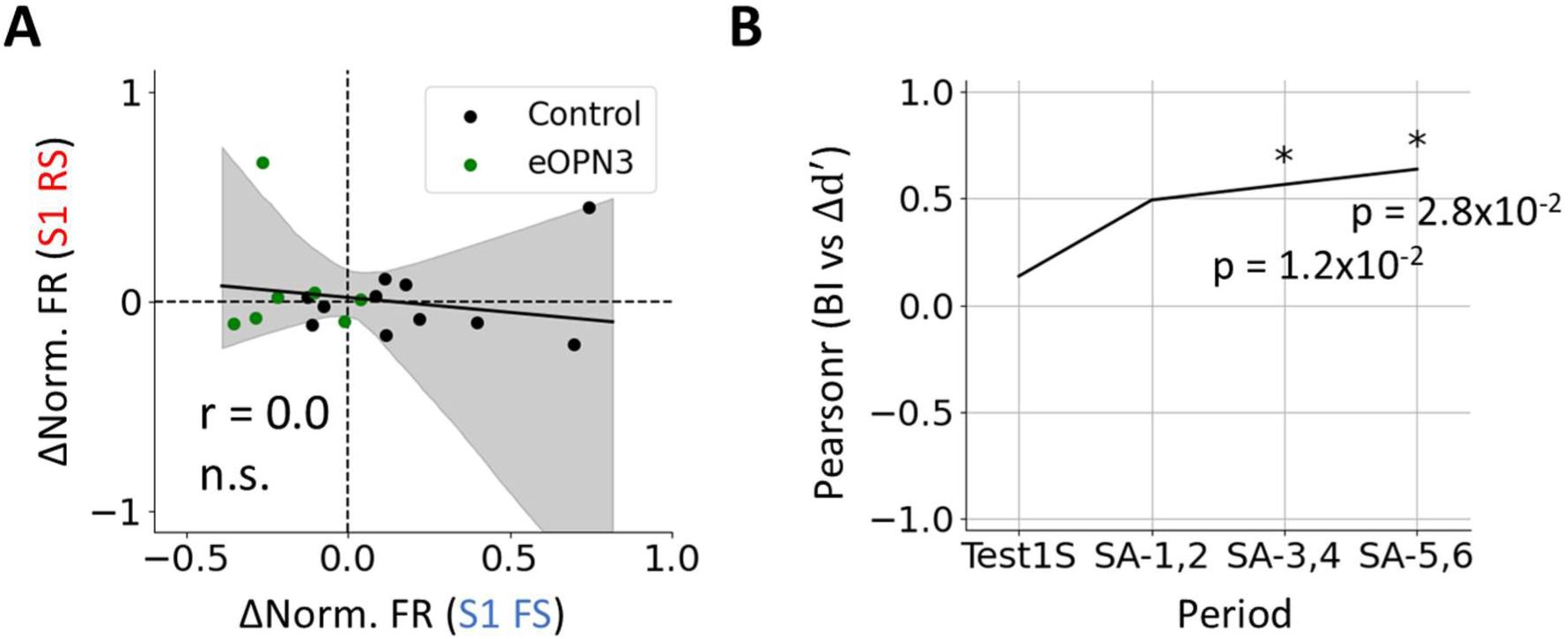
S1 FS controls S1 RS burst firing, but not firing rate, and facilitates blocking, related to Figure 4. A, The differences in S1 FS normalized firing rates between SA-1 and Test1 S within 300 ms after cue were plotted against the differences in S1 RS normalized firing rates between SA-1 and Test1 S within 300 ms after cue. A prominent correlation was not observed (one-sided bootstrap test). Black dots represent control mice, and green dots represent OPN3 mice. B, Correlation coefficients between burst index (BI) and Δ*d*^′^ across each period, related to Figure 4I. Positive correlation progressively increased during Phase2 and became significant in SA-3,4 and SA-5,6 (one-sided bootstrap test).

**Figure S7.**
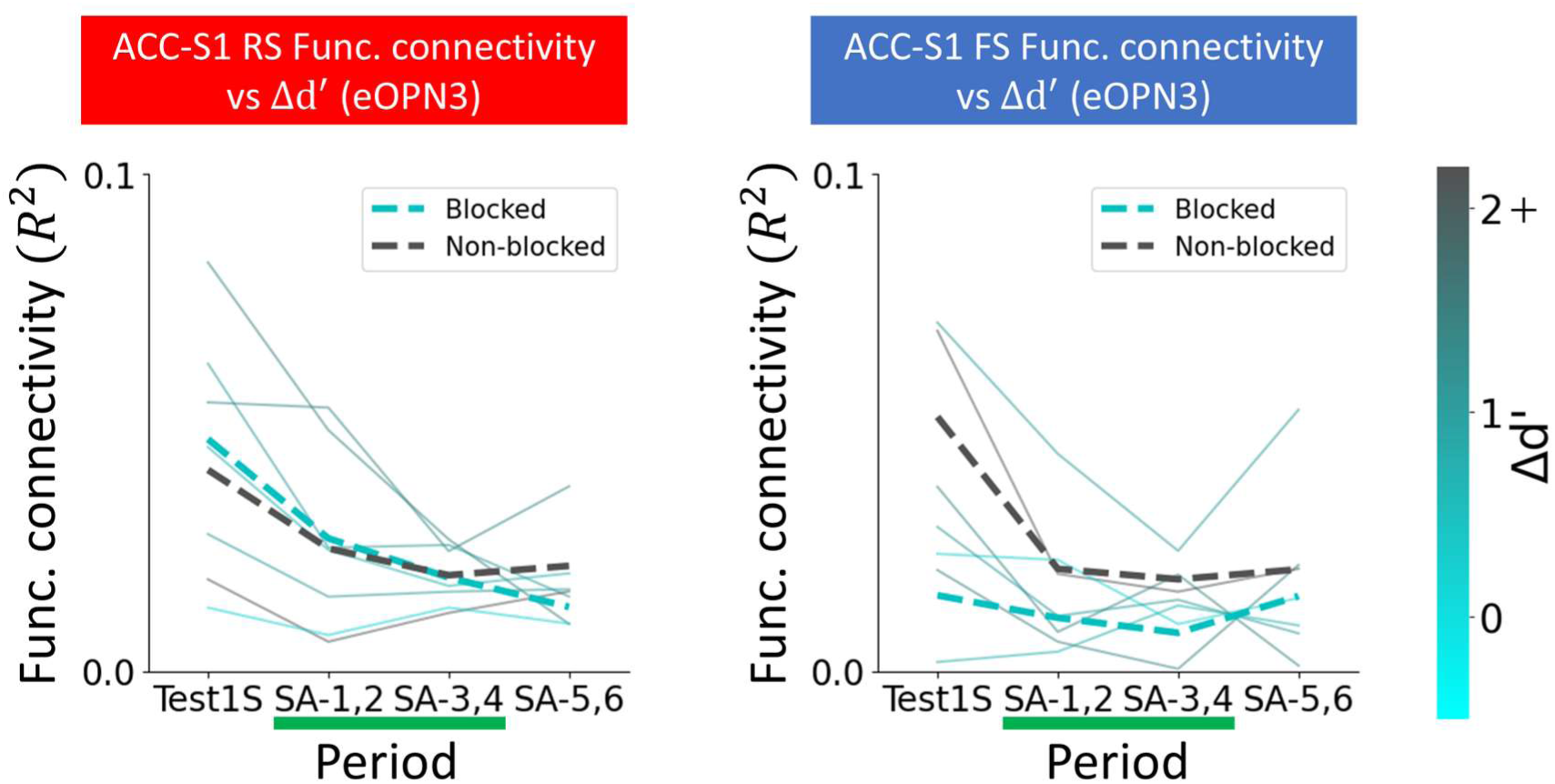
Suppression of the top-down input alters the relationship between the functional connectivity and blocking, related to Figure 5. Functional connectivity (*R*^2^) between ACC and S1 RS (Left) and between ACC and S1 FS (Right) of eOPN3 mice. Solid lines indicate individual animals colored by Δ*d*^′^ with the color scale capped at 2.1, and dashed lines indicate the group mean. Green bars represent light illumination periods. Unlike control mice (Figure 5C), the relationship between ACC–S1 FS functional connectivity and learning score was altered in eOPN3 mice. One possible interpretation is that mice with stronger ACC–S1 FS coupling were more strongly affected by the suppression of top-down inputs, resulting in a greater disruption of inhibitory influence on S1 and a greater tendency to become Non-blocked mice.

**Figure S8.**
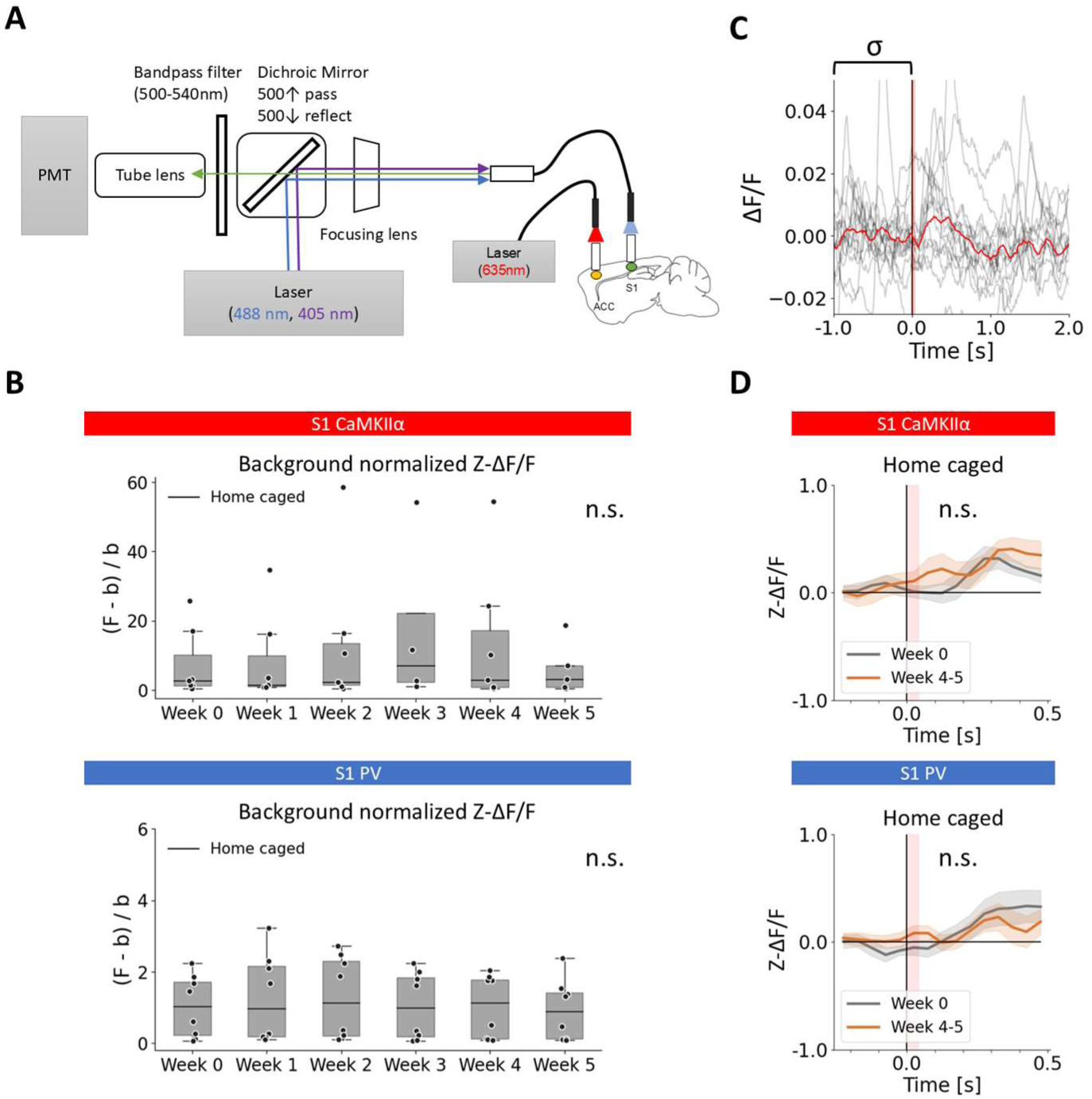
Photometry setup and home-caged control, related to Figure 6. A, Schematic of the fiber photometry setup. The 488 nm and 405 nm excitation lasers were reflected by a dichroic mirror (< 500 nm) and delivered to the S1 cannula through a focusing lens. Emitted GCaMP fluorescence was filtered with a dichroic mirror (> 500 nm) and a band-pass filter (500-540 nm) and detected by photomultiplier tube (PMT). A 635 nm red laser was connected to the ACC cannula to deliver optical stimulation to ACC. B, To validate the stability of GCaMP fluorescence in the photometry recordings, a home-caged control group was examined. Background-normalized GCaMP fluorescence was defined as (F − b)/b, where b represents autofluorescence of brain measured via the ACC cannula and F represents the resting-state GCaMP fluorescence measured through the S1 cannula. Week 0 was defined as the initial recording session, and recordings were performed once per week thereafter. Data from S1 excitatory (exc.) neurons (Top; n=7) and S1 PV neurons (Bottom; n=8). No significant week-dependent changes were observed in either group (one-way ANOVA). C, Representative S1 Ca^2+^ response after ACC activation in Week 0. GCaMP fluorescence (F) was processed as ΔF/F = (F − F_0_)/F_0_, where F_0_ denotes the mean fluorescence during the 1 s before activation. Black lines show individual trials, and the red line is the average. To facilitate comparisons across days, ΔF/F was normalized by the standard deviation of 1 s before activation (σ), yielding Z-scored ΔF/F (Z-ΔF/F = ΔF/F /σ). Red shaded areas indicate the ACC activation window (40 ms). D, S1 Ca^2+^ responses of home caged group in Week 0 and Week 4-5 (corresponding to before and after Phase1 in the blocking task, respectively). Red shaded areas indicate the ACC activation window (40 ms). Neither S1 exc. (Top) nor S1 PV neurons (Bottom) showed significant differences (one-sided paired bootstrap test with max-T correction).

**Figure S9.**
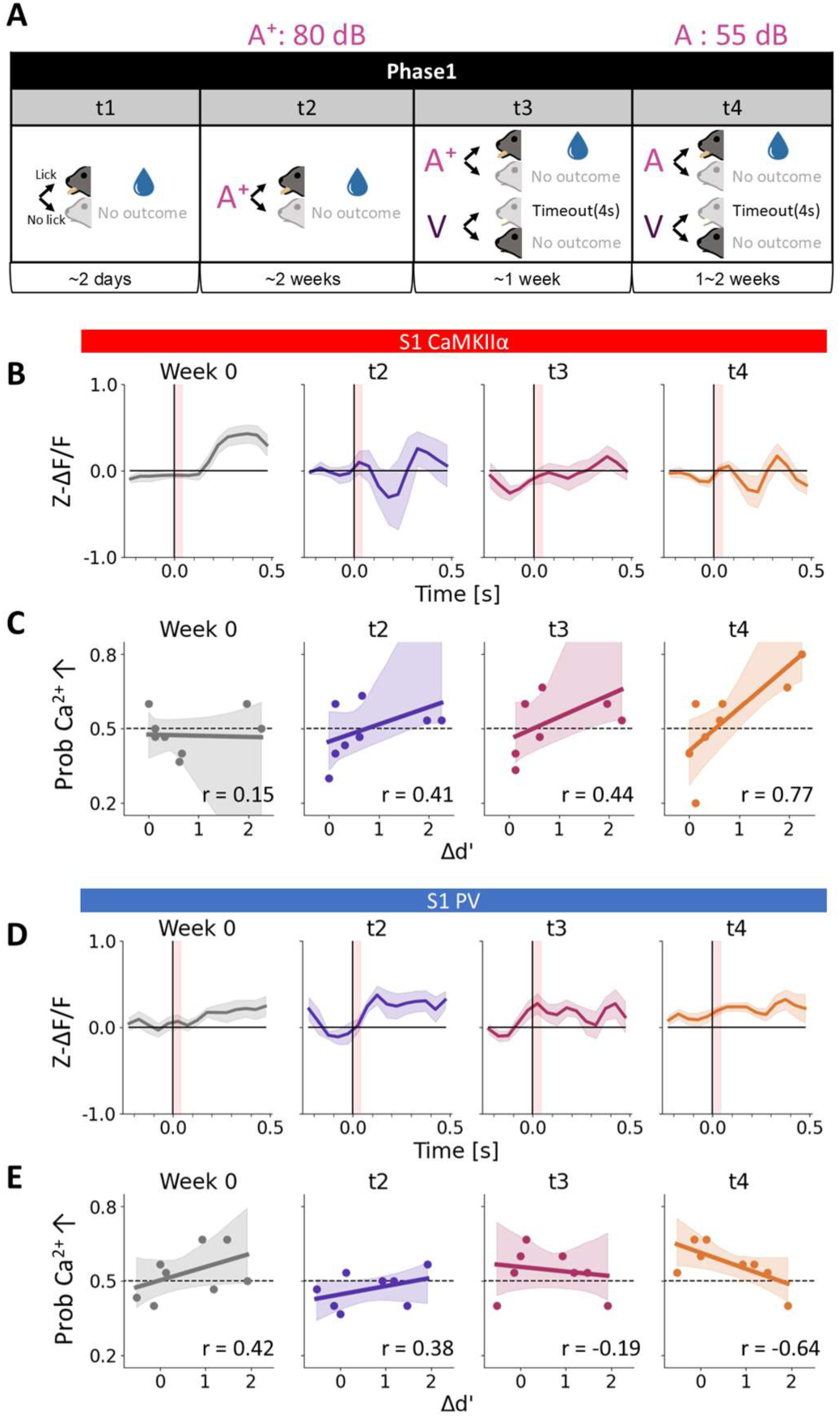
Gradual changes in S1 Ca^2+^ response to ACC activation across Phase1, related to Figure 6. A, Schematic of the four learning stages in Phase1. At t1, mice learned to lick for reward. At t2, mice performed a Go task in response to cue A of 80 dB (A⁺). At t3, mice performed a Go task in response to cue A⁺ and a No-go task in response to cue V. At t4, mice performed a Go task in response to cue A (55 dB) and a No-go task in response to cue V. B, ACC-activated S1 excitatory (exc.) neurons’ Ca^2+^ response at each learning stage in Phase1. Mean ± s.e.m. Red shaded areas indicate the ACC activation window (40ms). The data for Week 0 and t4 are the same as those shown in Figure 2E. C, For each learning stage in Phase1, probability of trials showing increased S1 exc. Ca^2+^ responses at 100-150 ms after ACC activation onset relative to 50 ms before the onset was plotted against the learning score (Δ*d*^′^). Mean regression (solid line) ± 95% CI (shaded). The data for Week 0 and t4 are the same as those shown in Figure 2G. The correlation coefficient gradually increases as Phase1 progresses. D, Same analysis as (B) for S1 PV neurons. The data for Week 0 and t4 are the same as those shown in Figure 2K. E, Same analysis as (C) for S1 PV neurons at 0-50 ms after ACC activation onset. The data for Week 0 and t4 are the same as those shown in Figure 2K. The correlation coefficient gradually decreases as Phase1 progresses.

## STAR★Methods

### KEY RESOURCES TABLE

| REAGENT OR RESOURCE | SOURCE | IDENTIFIER |
| --- | --- | --- |
| Antibodies |  |  |
| Chicken anti-GFP | Abcam | Cat #ab13970; RRID:AB_300798 |
| Rabbit anti-mCherry | Abcam | Cat #ab167453; RRID:AB_2571870 |
| Rabbit anti-Somatostatin | Peninsula | Cat #T-4103; RRID:AB_518614 |
| Mouse anti-Parvalbumin | Swant | Cat # 235; RRID:AB_10000343 |
| Alexa Fluor 488 goat anti-chicken IgG | Thermo Fisher Scientific | A11039; RRID:AB_2534096 |
| Alexa Fluor 555 F(ab')<sub>2</sub> goat anti-rabbit IgG | Thermo Fisher Scientific | A21430; RRID:AB_10374475 |
| Alexa Fluor 647 goat anti-mouse IgG | Jackson ImmunoResearch | Cat #111-607-185; RRID:AB_2632544 |
| Bacterial and virus strains |  |  |
| AAV2-CaMKIIα-mGFP-T2A-Synaptophysin-mRuby-WPRE | This paper | N/A |
| AAV1-hSyn-Cre-WPRE-hGH | Addgene | Cat #105553-AAV1 |
| AAV1-hDlx-flex-GFP-Fishell_6 | Addgene | Cat #83895-AAV1 |
| AAV1-CaMKII-eOPN3-mScarlet | Addgene | Cat #125712-AAV1 |
| AAV1-CaMKII-mCherry | Addgene | Cat #114469-AAV1 |
| AAV1-CAG-mCherry-2a-hM4D-nrxn1a-WPRE | This paper | N/A |
| AAV1-CAG-GFP | Addgene | Cat #37825-AAV1 |
| AAV8-hSyn-ChRmine-mScarlet-Kv2.1-WPRE | Addgene | Cat #130995-AAV8 |
| AAV1-CaMKIIα-jGCaMP8m-WPRE | Addgene | Cat #176751-AAV1 |
| AAV1-syn-FLEX-jGCaMP8m-WPRE | Addgene | Cat #162378-AAV1 |
| Chemicals, peptides, and recombinant proteins |  |  |
| CTB Alexa Fluor 647 conjugate | Thermo Fisher Science | Cat #C34778 |
| Hoechst 33342 | Nacalai tesque, Inc. | Cat #19172-51 |
| DAPI | Sigma | Cat #D9542 |
| Experimental models: Organisms/strains |  |  |
| Mouse: C57BL/6NcrSlc | Japan SLC |  |
| Mouse: PV-ires-cre | Jackson Laboratory | Strain #008069; RRID: [IMSR_JAX:008069](https://identifiers.org/IMSR_JAX:008069) |
| Recombinant DNA |  |  |
| pAAV-CaMKIIα-mGFP-T2A-Synaptophysin-mRuby-WPRE | This paper | N/A |
| pAAV-CAG-mCherry-2a-hM4D-nrxn1a-WPRE | This paper | N/A |
| Software and algorithms |  |  |
| Fiji | NIH | https://fiji.sc/ |
| Python 3.10 | The Python Software Foundation | https://www.python.org/ |
| Spikeinterface | Buccino et al.<sup>64</sup> | https://github.com/SpikeInterface/spikeinterface |
| Kilosort 2.5 | Pachitariu et al.<sup>65</sup> | https://doi.org/10.5281/ZENODO.4482749 |
| MATLAB | MathWorks | https://www.mathworks.com |
| Labview | National instruments | https://www.ni.com |
| Custom code | This paper | https://github.com/toppo365/proactive_filter.git |

### Animals

All experimental procedures were approved by the Animal Experiments Committees of the RIKEN and the National Institute for Physiological Sciences. Wild-type (WT) mice used in this study were male C57BL/6N mice purchased from Japan SLC, Inc. PV-Cre mice (JAX #008069) were purchased from Jackson Laboratory and maintained by in-house breeding, and heterozygous animals were used for all experiments. Mice were predominantly male; however, female mice were included in a subset of experiments that did not involve the behavioral task. All animals were housed in a standard plastic cage under a reversed 12-h light/dark cycle. Mice had ad libitum access to food throughout the experiment and to water except during the behavioral task.

### Blocking task

Mice were implanted with a custom-made metal head bar under isoflurane anesthesia and allowed to recover for at least three days before behavioral training. Water restriction was initiated one day before training and maintained throughout the task, with body weight kept above 80% of baseline. During task sessions, mice were head- and hindlimb-fixed, and licking responses were detected using a non-contact optical sensor and recorded with a data acquisition system (DAQ, National instruments). All the behavioral training program was operated using a custom written MATLAB code.

Behavioral training of Phase1 consisted of four stages (t1-t4, see Extended Data Figure 5a). In t1, mice were habituated to the task setup and trained to associate licking with reward; each lick triggered the reward delivery of about 2μL of 5% sucrose water via a lick port. It typically completes within two sessions. In t2, mice were trained on an auditory (A⁺) Go task. A salient auditory cue (A⁺) was delivered from a speaker positioned about 30 cm posterior to the left side of the mice and consisted of a 12 kHz square wave (duty cycle 0.5, 80 dB) of 150 ms duration. Licks detected within a response period of 1.2s following cue onset were defined as hits and were rewarded. After reward presentation, a delay period of 3 s was imposed for reward consumption. Before the next cue presentation, a no-lick period, randomly varied between 0.5 and 4 s, was introduced, and licks occurring during this period triggered an additional no-lick period (timeout). The duration of timeout manually increased from 1 to 4 s according to mice’s performance. Catch trials, in which no cue was presented, were interleaved to calculate false alarm (FA) rate. The proportion of catch trials increased from 25% to 50% over training. Mice that achieved a hit rate ≥90% and a FA rate ≤25% under the final conditions (timeout = 4 s, catch trials = 50%) were advanced to the next training stage. In t3, mice were trained on an auditory (A⁺) Go / visual (V) No-go task. The salient auditory cue (A⁺) was identical to that used in t2, whereas the visual cue (V) consisted of a flashing light (10 Hz, duty cycle 0.5) presented for 400 ms from a light source positioned ∼3 cm in front of the face. Hits for cue V triggered a 4 s timeout. Cue A⁺ and V were presented with equal probability (1:1) in a randomized order. A no-lick period was randomly varied between 0.5 and 4 s, and violations triggered a 4 s timeout. The proportion of catch trials was set to 14.3%. Mice that achieved a hit rate for cue A⁺ ≥90% and both a hit rate for cue V and FA rate ≤25% in at least two sessions were advanced to the next training stage. In t4, mice were trained on the same task structure as in t3, except that the sound intensity of the auditory cue was gradually reduced from 80 dB (A⁺) to 55 dB (A) across training. For mice used for electrophysiological recordings, to minimize behavioral changes induced by electrodes insertion, mice were exposed to bright white light for around 30 min under head- and hindlimb-fixed conditions before the initiation of t4 training. Using the 55 dB auditory cue (A), mice that achieved a hit rate for cue A ≥90% and both a hit rate for cue V and FA rate ≤25% for at least three sessions were considered to have completed Phase1. Mice that completed Phase1 underwent hair removal from the left dorsocaudal trunk, and the blocking test was performed on the following day.

On the day of the blocking test, mice were head- and hindlimb-fixed, and around 20 trials of t4 were first conducted to stabilize the performance. The blocking test consisted of three consecutive phases: Test1, Phase2, and Test2. Test1 and Test2 were auditory (A) Go / somatosensory (S) No-go tasks with the same structure as t4. Cue A was identical to that used in t4, whereas the somatosensory cue (S) was delivered via adhesive surface electrodes (Skintact electrode, Leonhard Lang GmbH) cut to around 0.5 cm width and attached to the left dorsocaudal trunk, through which a 3 mA current pulse of 0.1 ms duration was applied. The order of cue A and S presentation (18 times each) was determined using an identical pseudorandom sequence. Phase2 were a compound auditory–somatosensory (SA) Go / visual (V) No-go task, with the same structure as t4. Cue SA and V were each presented 117 times in an identical pseudorandom order. Only mice with a hit rate ≥ 70% for cue SA in Phase2 were included in the analysis.

### Control (overshadowing) task

For control experiment, overshadowing task was performed to assess experience-independent learning performance. In this experiment, Phase1 training was conducted using an alternative auditory cue (A’), consisted of a 7 kHz square wave (duty cycle 0.5, 80 dB) for 150 ms duration presented from a speaker positioned around 5 cm anterior to the right side of the face. Using this auditory cue (A’), mice underwent the same training procedures as the blocking task in t1–t3, and mice that achieved a criteria where a hit rate for cue A’ ≥90% and both the hit rate for cue V and FA rate ≤25% for at least five sessions in t3 were considered to have completed Phase1. During the blocking test, Test1 was performed using cue A’, whereas Phase2 and Test2 were performed using cue A. Because many mice initially failed to respond to the novel compound cue (SA) at the beginning of Phase2, trials in which mice did not lick within 0.5 s after cue SA onset were followed by automatic reward delivery. Only animals with a hit rate ≥ 50% for cue SA in Phase 2 were included in the analysis.

### Learning score

To quantify reward expectations for cue S in Test1 and Test2, we used the discrimination index *d*^′66^. *d*^′^ was defined using the inverse of the standard normal cumulative distribution function, *Z*, as

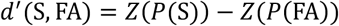

where *P*(S) is the hit rate for cue S and *P*(FA) is FA rate in each test phase. To minimize the influence of extinction learning due to the absence of reward following the lick responses after cue S, *P*(S) was calculated using only the first eight presentations of cue S. To quantify changes in reward expectations for cue S across Phase2, a learning score (Δ*d*^′^) was defined as the difference in *d*^′^ between Test1 and Test2,

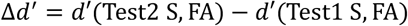

To minimize the contribution of extinction learning of cue S in Test1 to subsequent associative learning in Phase2, mice with either a high lick probability to Test1 S (*P*(Test1 S) > 0.5) or a high reward prediction to Test1 S (*d*′(Test1 S, FA) ≥ 1) were excluded from the analysis.

### Stereotaxic Injections

Mice aged 6-12 weeks were anesthetized with either 2% isoflurane or a mixture of medetomidine, midazolam, and butorphanol in saline (0.05 mg kg^−1^, 5 mg kg^−1^ and 0.5 mg kg^−1^ respectively, injected intraperitoneally), and positioned in a stereotaxic holder. After scalp incision and exposure of the skull, target coordinates were determined. Target coordinates relative to bregma (β) were as follows: S1Tr, anterior–posterior (AP) −1.4 mm, medial–lateral (ML) +1.5 mm; ACC, AP +2.2 mm, ML +0.35 mm. Small craniotomies were made at the corresponding locations using a dental drill. Adeno-associated virus (AAV) vectors or cholera toxin subunit B (CTB) were delivered using glass micropipettes connected to a nanoliter injector (Picospritzer, General Valve; Nanoject, Drummond). Injections were performed perpendicular to the cortical surface at depths spanning from 650 µm to 250 µm in S1Tr and from 700 to 600 µm in ACC, with each injection delivered over ≥15 min. For in vivo photometry experiments, injections into S1Tr were made 0.4 mm posterior to the target coordinate and angled at 30°(posterior-to-anterior) to avoid tissue damage at the recording site.

### Anatomical tracing of top-down inputs to S1Tr

To identify dFC regions projecting to S1Tr, 200 nL of 0.5% CTB Alexa Fluor 647 conjugate (Thermo Fisher C34778) was injected into S1Tr. To visualize axon and presynaptic terminals projecting from ACC to S1Tr, AAV2-CaMKIIα-mGFP-T2A-Synaptophysin-mRuby-WPRE (1.1×10¹² GC/mL, 200nL), generated by combining plasmids and packaging it into AAV, was injected into ACC.

To label S1Tr inhibitory neurons receiving direct projections from ACC, we followed the previous research^42^. Briefly, AAV1-hSyn-Cre-WPRE-hGH (1.4×10¹³ GC/mL, 200nL, Addgene 105553-AAV1) supplemented with Hoechst 33342 solution (0.1 mg/mL) was injected into ACC, and AAV1-hDlx-flex-GFP-Fishell_6.^67^ (5.2×10¹² GC/mL, 200nL, Addgene 83895-AAV1) was injected into S1Tr.

At the end of the experiments, mice were transcardially perfused with 4% paraformaldehyde (PFA) in phosphate-buffered saline (PBS), and brains were post-fixed in 4% PFA in PBS at 4℃. For CTB injections, perfusion and fixation were performed 7 days after injection. For AAV injections, perfusion and fixation were performed 3–4 weeks after injection. After post-fixation, brains were washed in PBS and sectioned at 50 µm thickness either with a vibrating microtome or with a freezing microtome after cryoprotection in PBS containing 20% sucrose at 4℃ for 1–3 days. Sections were subjected to 4’,6-diamidino-2-phenylindole (DAPI) staining or immunohistochemistry as required, mounted on glass slides, and imaged using a fluorescence or laser-scanning confocal microscope. For DAPI staining, sections were incubated with DAPI (1 µg/mL, Sigma-Aldrich D9542) for 10–15 min and washed with PBS. For immunohistochemistry, sections were incubated in a blocking solution consisting of PBS containing 0.5% Triton X-100 and 5% normal goat serum for 30 min and then incubated overnight at 4℃ in the blocking solution containing primary antibodies against GFP (chicken, Abcam ab13970; 1:1000), RFP (rabbit, Abcam ab167453; 1:1000), somatostatin (rabbit, Peninsula T4103; 1:1000), and parvalbumin (mouse, Swant 235; 1:1000). After washing with PBS, sections were incubated for 2 h at room temperature with the appropriate fluorophore-conjugated secondary antibodies (Alexa Fluor 488–conjugated goat anti-chicken IgG, Invitrogen A11039; Alexa Fluor 555 F(ab’)₂ fragment goat anti-rabbit IgG, Invitrogen A21430; Alexa Fluor 647–conjugated goat anti-mouse IgG, Jackson ImmunoResearch 111-607-185; all 1:500), with NeuroTrace Blue (Invitrogen N21479) at a 1:100 dilution or DAPI (1 µg/mL, Sigma-Aldrich D9542) for whole-cell labeling, followed by washing with PBS.

### Top-down input suppression experiments

For DREADD-mediated suppression, WT mice aged 8 weeks received injections of AAV1-CAG-mCherry-2A-hM4D-nrxn1a-WPRE (1.0×10¹² GC/mL, 200 nL, this paper) or the control virus AAV1-CAG-GFP (1.0×10¹² GC/mL, 200 nL, Addgene 37825-AAV1) into ACC. For optogenetic suppression, WT mice aged 8 weeks received injections of AAV1-CaMKII-eOPN3-mScarlet^68^ (1.95×10¹² GC/mL, 200nL, Addgene 125712-AAV1) or the control virus AAV1-CaMKII-mCherry (1.4×10¹² GC/mL, 200nL, Addgene 114469-AAV1) into ACC. After head-bar implantation and recovery, mice were subjected to water restriction and trained on the blocking task. On the day of the blocking test, typically 4-5 weeks after viral injection, mice were lightly anesthetized with isoflurane (∼0.5%) and head-fixed, after which a craniotomy was performed over the S1Tr and the cortical surface was covered with silicone adhesive (Kwik-Sil, World Precision Instruments). More than 30 minutes later, allowing sufficient recovery from anesthesia, mice were head- and hindlimb-fixed, the silicone adhesive was removed, and an optical fiber was positioned over the S1Tr cortical surface. The blocking test was conducted, and 535 nm laser light (Shanghai Laser & Optics Century Co., Ltd.) adjusted 10 mW at the cortical surface was delivered through the optical fiber as a 500 ms light pulse during the first two-thirds of catch trials in Phase2.

### In vivo electrophysiological recording

Neuronal activity during the blocking test was recorded using an Open Ephys acquisition system. For recordings in S1Tr, a 4-shank 64-channel probe (A4×4-tet-5mm-150-200-121, NeuroNexus Technologies, Inc.) was used, whereas recordings in ACC were performed using a 6-shank 64-channel probe (F64, tip-sharpened, Cambridge Neurotech, Ltd.). On the day of the blocking test, mice were lightly anesthetized with isoflurane (∼0.5%), and craniotomy and dura removal were performed over S1Tr and ACC. Anesthesia was then discontinued, and mice were head- and hindlimb-fixed at the task environment under bright white illumination. (Mice had been habituated to this light illumination (see Blocking task section)). To enable post hoc tracking of electrode positions, probes were coated with the infrared fluorescent dye (DiD) before insertion. Probes were inserted into ACC and then S1Tr to a target depth of approximately 1,000 µm from the cortical surface, primarily sampling activity from layers 5 and 6, as judged by depth and changes in firing rates. Probe insertion was performed at a rate of approximately 100 µm per minute. When combined with optogenetic suppression of top-down inputs with eOPN3, an optical fiber was positioned as close as possible to the S1Tr cortical surface and the silicon probe. 1% agarose dissolved in Ringer’s solution was applied to the cortical surface to prevent tissue drying and probe movement. A common ground and reference electrode were placed in the agarose. After waiting for at least 15 min following probe insertion, the white illumination was turned off, and the blocking test and neuronal recordings sampled at 30 kHz were initiated. After completion of the blocking test, mice were perfused and histological analyses were performed to confirm the probe positions.

### Processing of neuronal data

Recorded signals were band-pass filtered between 600 and 6,000 Hz and processed with median-based common average referencing separately for each probe in ACC and S1Tr. To avoid contamination by electrical artifacts induced by somatosensory stimulation, data within ±2 ms around the electrical stimulus onset were excluded from analysis. Spike sorting was performed automatically using Kilosort2.5^65^, followed by manual inspection to verify unit identity. Units with an inter-spike interval (ISI) violation rate below 0.5 were classified as single units. Units that exhibited complete loss of spiking activity during the recording session were manually excluded. When multiple spikes were detected within a 1 ms window, they were regarded as double-counted spikes and counted as a single spike. Mice from which at least 10 units were recorded in both ACC and S1Tr were included in the analysis. Units were further classified based on waveform shape: units with a peak-to-valley duration shorter than 0.5 ms were classified as fast-spiking (FS) units, whereas those with a duration of 0.5 ms or longer were classified as regular-spiking (RS) units. For calculating normalized firing rates, firing rates of each unit were divided by the mean population firing rate of the corresponding unit class (S1 RS, S1 FS, ACC RS, or ACC FS) during the first 300 ms after cue onset in Test1 S.

### Burst index

Burst firing was defined as spikes with an inter-spike interval (ISI) of <5 ms^58,69^. To quantify the increase in burst firing after cue compared with baseline, we calculated a burst index (BI) for each unit as a difference in burst fraction^41^:

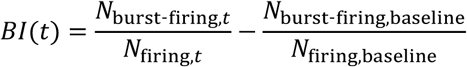

where N denotes the number of total firing events (firing) or burst-firing events (burst-firing) within a given time window. The baseline period was defined as 300–100 ms before cue onset.

### Partial least squares

To estimate functional connectivity between ACC and S1, partial least squares (PLS) analysis was applied. Spike trains from ACC and S1 were converted to firing rates using 20 ms bins from 100 ms before to 600 ms after each cue onset (Test1 S, SA-1,2, SA-3,4 and SA-5,6). To isolate trial-by-time covariations in neuronal activity removing shared cue-locked components in ACC and S1, the trial-averaged firing rates were subtracted from each unit, analogous to the computation of noise correlations. Since our goal was to quantify how well ACC activity predicts S1 population activity, PCA was applied separately to S1 RS and S1 FS populations, and the first principal component (PC1) was used for low dimensional S1 population activity. This procedure yielded a robust estimate of S1 population activity while minimizing variability arising from differences in recorded units across animals. ACC and S1 activity in each period were then concatenated across time bins to form activity matrices ℝ^nbin×nunit^, where *n*_bin_ denotes the total number of time bins and *n*_unit_ denotes the number of units, which varied across animals. Since we used PC1 activity for S1 RS and FS, *n*_unit_ = 2 for S1 in all mice.

Resulting ACC and S1 activity (*X* and *Y*, respectively) were linearly projected onto 2-dementinal latent space ( *XU* and *YV*, respectively), using projection matrices (*U* ∈ ℝ^nunit×2^ and *V* ∈ ℝ^2×2^, respectively). The projection matrices were optimized to maximize the covariance between the projected ACC and S1 latent representations as

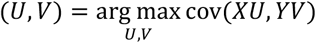

Because *V* is a square matrix and its inverse can be computed, the functional connectivity matrix between ACC and S1 (*W*) was defined as *W* = *UV*^-1^. Functional connectivity was quantified using the coefficient of determination,

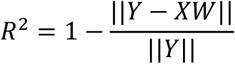

which measures the extent to which trial-by-time fluctuations in S1 population activity (Y) can be linearly explained by corresponding fluctuations in ACC activity (X), thereby reflecting the strength of their functional connectivity.

### In vivo Fiber Photometry

To examine ACC-activated Ca^2+^ responses of excitatory neurons in S1Tr, WT mice aged 6 weeks received injections of AAV8-hSyn-ChRmine-mScarlet-Kv2.1-WPRE^43^ (1×10¹² GC/mL, 200 nL, Addgene 130995-AAV8) into ACC and AAV1-CaMKIIα-jGCaMP8m-WPRE (1.8×10¹³ GC/mL, 200 nL, Addgene 176751-AAV1) into S1Tr. To examine ACC-activated Ca^2+^ responses of PV-positive neurons in S1Tr, PV-Cre mice aged 6 weeks received injections of the same virus as above in ACC and AAV1-syn-FLEX-jGCaMP8m-WPRE^70^ (2.0×10¹³ GC/mL, 200 nL, Addgene 162378-AAV1) into S1Tr. At 8 weeks of age, head bars were implanted (see Blocking task section). At 9 weeks of age, optical cannulas (0.5 numerical aperture, 400 µm core diameter) were implanted over the cortical surfaces of ACC and S1Tr. Craniotomies were performed over the target regions, the dura mater was carefully removed, and the optical cannulas were placed in contact with the cortical surfaces. The exposed brain surface was covered with 1% agarose dissolved in saline, protected with silicone adhesives (Kwik-Sil, World Precision Instruments), and the cannulas were then attached to the skull using dental cement mixed with black pigment, blocking ambient light. Fiber photometry recordings were initiated after a recovery period of at least 3 days following cannula implantation.

For fiber photometry recordings, ACC activation was conducted using a 635 nm red laser (Shanghai Laser & Optics Century Co., Ltd.) adjusted to an intensity of around 10 mW at the cortical surface. S1Tr GCaMP excitation was performed using a 488 nm laser, together with a 405 nm laser (Coherent Corp.) to remove artifacts related hemodynamic and motion. Both excitation lasers (405, 488 nm) were adjusted to around 100 µW at the cortical surface and temporally interleaved by alternating 4 ms laser on and 6 ms off periods. Fluorescence collected through the optical cannula was band-pass filtered (500–540 nm; FBH520-40, Thorlabs) to isolate GCaMP8m emission, projected onto photodetector (PMTSS, Thorlabs), amplified via a current-to-voltage converter (TIA60, Thorlabs), and digitized using a data acquisition system (DAQ, National Instruments) (see Extended Data Figure 4a). Signals were sampled at 5,000 Hz and downsampled to 100 Hz by averaging voltage values during laser-on periods for each excitation wavelength. Mice were randomly assigned to either blocking task group or home-caged group and underwent photometry recordings once per week for five consecutive weeks to measure ACC-activated Ca^2+^ responses in S1Tr. During recordings, mice were head- and hindlimb-fixed, baseline autofluorescence was first measured through the ACC-side cannula, and then GCaMP fluorescence was recorded through the S1Tr-side cannula. For ACC activation, photostimulation with the 635 nm laser was delivered through the ACC-side cannula. The photostimulation consisted of repeated cycles of 2 ms laser on followed by 3 ms off periods, with stimulus durations pseudo-randomly varied across trials (20–640 ms); unless otherwise stated, data are shown for the 40 ms condition. Each stimulus was repeated 15 times with an inter-stimulus interval of 7 s.

Recorded signals (405, 488 nm) from the ACC and S1Tr were separately filtered with a 30 s median filter to correct for photobleaching, and the 405 nm signal was linearly regressed onto the 488 nm signal to remove artifacts. The residual signals from the ACC-side and S1Tr-side cannula were considered as baseline autofluorescence (b) and GCaMP fluorescence (F), respectively. To estimate GCaMP fluorescence relative to baseline autofluorescence, we calculated (F − b)/b. To quantify ACC-activated Ca^2+^ responses, fluorescence changes were calculated as ΔF/F = (F − F₀)/F₀, where F₀ is the mean fluorescence (F) of 1 s before the activation onset. To further remove day-to-day and inter-animal variability, ΔF/F values were Z-scored (Z-ΔF/F) by dividing by the standard deviation of the fluorescence (F) of 1 s before the activation onset. Mice showing no detectable S1Tr responses to ACC photostimulation or exhibiting a marked decrease in estimated GCaMP fluorescence ((F− b)/b) over the 5-week recording period were excluded from analysis. After completion of recordings, mice were perfused and histological analyses were performed; mice with insufficient viral expression in the ACC or S1Tr, or with evident brain damage, were also excluded.

## Statistical analysis

Statistical analyses are described in the figure legends. Nonparametric tests were used throughout: Wilcoxon rank-sum tests for two-group comparisons, Wilcoxon signed-rank tests for paired comparisons, permutation test with max-statistic correction across time for assessing the temporal extent of group differences, bootstrap tests for correlations, two-sample permutation tests for group comparisons of correlations, and paired bootstrap tests with max-T correction for time-series analyses.

To assess the temporal extent of group differences following cue onset, we performed a one-sided permutation test with max-statistic correction across time. Neural activity was first averaged within consecutive 100-ms bins. For each time point *x*, the test statistic was defined as the difference between groups in the cumulative mean response from cue onset to bin *x*. Group labels were randomly permuted 5,000 times, and the maximum statistic across all time points was retained for each permutation to construct the null distribution. One-sided p values were calculated from this max-statistic distribution.

For correlation analyses, data were resampled with replacement 5,000 times to generate an empirical null distribution of the correlation coefficient; one-sided P values were calculated as the proportion of bootstrap samples exceeding the observed coefficient, and 95% confidence intervals (CI) were estimated from the bootstrap distribution.

For group comparisons of correlations, group labels were randomly permuted 5,000 times, and the difference in correlation coefficients between groups was recalculated for each permutation to generate an empirical null distribution. One-sided P values were computed as the proportion of permuted differences exceeding the observed difference.

For time-series analyses in fiber photometry experiments, Ca²⁺ responses were analyzed in 50 ms bins, and differences between before and after Phase1 were calculated for each mice within the 300 ms window following ACC activation onset. Under the null hypothesis of no mean difference, data were centered and resampled with replacement 5,000 times to generate an empirical null distribution. Corrected P values for each time bin were obtained using a max-T procedure by comparing each observed difference with the distribution of the maximum bootstrap statistic across all time bins. Statistical significance was defined as P < 0.05.

